# Thermodynamic Continuum Tumour Modelling Recovers Multicompartment Spheroid Structure and Supports Drug-Response Screening

**DOI:** 10.64898/2026.04.08.717345

**Authors:** Riley J. McNamara, Gloria M. Monsalve-Bravo, Sandra R. Stein, Glenn D. Francis, Mark C. Allenby

## Abstract

Patient-derived tumour spheroids provide experimentally tractable models of tumour growth and treatment response, but standard radius-based measurements do not directly identify the tissue-scale mechanisms that shape internal structure. This study develops and calibrates a thermodynamic continuum model of untreated spheroid internal structural growth, in which viable and necrotic cell populations, nutrient transport, mechanical relaxation, and cell-fate transitions are represented as coupled spatial fields. Because the model does not prescribe internal concentric spheroid compartments like traditional spheroid models, the experimentally reported intermediate (inhibited) radius was treated as an exploratory model-to-data mapping and optimised over different candidate field-derived radii. The calibrated model reproduced the experimental trajectories with aggregate uncertainty-weighted root-mean-square error (WRMSE) values ranging from 0.49 to 1.24 on the metric defined in the manuscript, and improved average fit quality by approximately 40% relative to an existing Greenspan-type compartment model. The model recovered the growth-inhibited and necrotic structure observed in spheroids as an emergent consequence of spatial cell-state dynamics rather than prescribing concentric compartments. Biological and nutrient-associated model parameters were locally constrained to the experimental data, although compensatory relationships were observed among nutrient-response and death parameters. The framework was further extended by introducing a drug transport field as a proof-of-concept demonstration of spatial treatment-response modelling. These results show that a continuum cell-growth model can recover classical spheroid structure while preserving flexibility for exploratory field-based analysis of drug response.

## 1 Introduction

Three-dimensional patient-derived tumour spheroids (PDTSs) have emerged as a central experimental platform in modern tissue engineering and precision oncology, and useful to inform the biophysical mechanisms governing tissue growth and behaviour [1]. These three-dimensional engineered systems preserve key features of host tumour architecture, cellular heterogeneity, environmental response, and even gene expression [2, 3]. Because of this, PDTSs have already been used in clinical settings to study tumour progression, interrogate therapeutic sensitivity, and inform personalised treatment decisions [4, 5]. Extending beyond clinical applications, these systems are also widely used for high-throughput drug screening in pharmaceutical research [6, 7, 8, 9, 10]. Their value lies in the fact that they preserve a spatially structured tumour-like environment outside the host tissue, while remaining experimentally tractable [11, 9, 12, 13, 14, 15, 16, 8, 17, 3].

Current drug-treated PDTS assays can report whether a spheroid becomes larger, smaller, more viable, or more necrotic under treatment, but these summary measurements do not by themselves explain which underlying biological mechanisms were impacted. This is a major limitation for precision oncology, where the useful question is not only whether a treatment changes the final spheroid size, but whether that response reflects altered proliferation, nutrient limitation, drug penetration, cell death, or some combination of these effects. Despite the clinical promise of PDTSs, sample availability can be variable [16, 3, 18], and experimental outcomes can vary greatly [13, 18]. There remains uncertainty in translating spheroid outcomes into clinical insight [5, 13]. Mechanistic mathematical models offer a route to bridge this gap by providing explicit representations of growth, death, nutrient transport, and tissue mechanics [19, 20, 12].

Existing approaches generally represent spheroids through empirical models, Greenspan-type models, agent-based models (ABMs), or continuum phase-field formulations [20, 21, 22]. Empirical models such as Gompertz, Logistic, or Fisher-KPP models generally model tumour volume or density directly. These models have been successful for curve fitting, but their empirical nature lies at the core of the clinical problem: they are limited in explaining which tissue-scale mechanisms have changed under treatment. ABMs model individual cells directly, and as a result resolve single-cell behaviour and stochastic fate decisions, but can become computationally expensive for multicellular systems [23, 24, 25, 26, 27]. Continuum models, often diffuse-interface or phase-field models, represent tissue, nutrient, and cell-state variables as coupled fields governed by partial differential equations [28, 27, 29, 30]. In this setting, internal structure can emerge from the governing transport and reaction dynamics, and model parameters can be interpreted as *effective* coarse-grained parameters that summarise many microscopic processes [31, 32, 30].

Within this modelling landscape, the canonical Greenspan model occupies a central historical and biological role [33]. A central organising concept in spheroid biology is the emergence of distinct radial compartments: an outer proliferative shell, an intermediate quiescent or growth-inhibited region, and a necrotic core [33]. This compartmental structure arises from the balance between nutrient diffusion, cellular consumption, and oxygen-dependent proliferation and death [34, 35, 36]. Dynamically, this is described as three distinct growth phases that a spheroid undergoes [33]. Phase 1 corresponds to initial growth of a fully viable spheroid, phase 2 to the emergence of the inhibited region, and phase 3 to the emergence of the necrotic core. Together these are known as the classical growth phases of spheroid growth [33, 37, 38]. The canonical Greenspan model and its descendants model these systems through radial compartments directly [39, 37]. This has been extremely useful for untreated spheroid growth, because the model structure mirrors the classical biological interpretation.

However, this same feature becomes a limitation when the intended application is drug response. Therapeutic agents penetrate spheroids through diffusion, creating spatially graded drug concentrations that couple to the same transport and consumption dynamics that govern nutrient distribution [40, 41]. In the same way that these three-dimensional systems model a diffusive continuum of nutrient, drug response is also inherently spatially dependent: outer cells experience higher exposure, while inner cells experience lower exposure. Even in the simple case where drug-induced death increases with local drug concentration, the strongest treatment effect may occur near the spheroid boundary, producing an outer region of drug-induced cell death rather than a stronger necrotic population within the classical necrotic core. This disrupts the traditional layered view of spheroids, where a fixed proliferative–inhibited–necrotic ordering is assumed. A model intended for drug-response analysis must therefore satisfy two requirements at once. It must recover the canonical compartment structure under untreated conditions, because that structure is a real and useful feature of spheroid biology. But it must not hard-code those compartments as the model geometry, because treatment can generate spatial patterns that do not respect the classical ordering.

This distinction also matters experimentally. Imaging and assay advances have expanded our ability to image these interior regions through high content brightfield, histological, and confocal microscopy [42, 43, 44, 45]. However, the sharp-boundary representation is an idealisation: in reality, cell viability, proliferation, and metabolic activity vary continuously with local nutrient concentration, and the transitions between compartments are gradual rather than discrete [46]. Furthermore, the definition of the intermediate region is ambiguous. Experiments have physically identified different intermediate regions that have been attributed as canonical structure [37, 45], but in reality this requires mapping an arbitrary thresholded experimental observable onto an ill-defined biological compartment. For a model to support treatment-response interpretation, these boundaries should therefore be treated as observables extracted from the simulated system, rather than as fixed states imposed by the model.

Continuum phase-field models therefore offer a route to resolving the modelling problem that motivates this paper. Cell populations are represented as spatially continuous volume fractions governed by reaction–diffusion mechanics, and internal structure emerges from coupled nutrient-growth dynamics rather than being prescribed [47, 48]. The model should be able to recapitulate the canonical proliferative–quiescent–necrotic architecture under untreated conditions, yet remain free to represent more complex spatial responses once a drug field is introduced. Furthermore, model parameters carry mechanistic interpretation as effective tissue-scale properties, such as effective growth and death rates, nutrient thresholds, and nutrient sensitivities, directly addressing the clinical need for biological reasoning beyond endpoint summary statistics. Any definition of compartment radii becomes a post-processing choice applied to the simulated fields, rather than a model constraint.

In this paper, we construct a continuum phase-field tumour growth model (CTM) to represent PDTS growth and calibrate it against untreated longitudinal spheroid imaging across multiple cell lines and seeding densities. The experimental dataset chosen characterises a particular observable as the classical boundary of the intermediate region. Within this paper, the mapping between model observables and the experimentally identified intermediate region of Browning *et al*. [37] is treated as an exploratory model-internal comparison. We recognise that the definition of this region is ambiguous, and that the observable Browning attributes to this intermediate boundary does not map directly onto a simulation observable. Therefore, we compare multiple candidate observable definitions within the same calibration workflow and identify which candidate gives the strongest relative agreement under the CTM. This result is conditional on the CTM being a useful working model; it does not establish the biological identity of Browning’s experimental boundary.

We further present an implementation of a drug field to simulate drug-treated conditions and demonstrate the model’s capacity for extension. We analyse varying drug strength, penetration, and mechanism of effect by modifying the untreated model’s cell-fate pathway dynamics. By establishing a continuum framework that can be calibrated to untreated spheroid data and recover the structural features identified by compartment-based models, we demonstrate that the same model infrastructure can support both descriptive and mechanistic analysis. We show how the reaction–diffusion equations governing nutrient transport can be extended to therapeutic agents while retaining the underlying cell-field representation.

Beyond drug screening, the same equations can apply to other experimental configurations, including spheroid formation, fusion assays, and scaffolded geometries, simply by changing initial and boundary conditions. This grants access to physical quantities that are difficult to measure directly, such as effective cell–cell interaction strengths and collective cell motility [47, 28]. The computational infrastructure built around the mathematical model supports Bayesian calibration, parameter sweeps, and integration into experimental workflows. Together, this study positions the CTM as a practical, transferable, and mechanistically grounded tool for analysing experimental spheroids and supporting future model-informed experimental design.

## 2 Theory – Mathematical Framework

We model a patient-derived tumour spheroid using a continuum phase-field framework adapted from Wise et al. [47]. The spheroid is described by volume fraction fields of viable and necrotic cell populations (*φ*_*V*_ , *φ*_*N*_), defined on a fixed domain Ω ⊂ ℝ^2^ subject to the constraint:

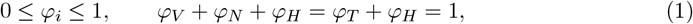

where *φ*_*T*_ = *φ*_*V*_ + *φ*_*N*_ is the total cell volume fraction and *φ*_*H*_ is the surrounding host medium.

### 2.1 Governing equations

Each cell population evolves according to a reaction–advection–diffusion equation:

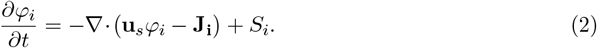

The diffusive Cahn–Hilliard-type flux **J**_**i**_ arises from the Helmholtz free energy and thermodynamic mixing theory [47]:

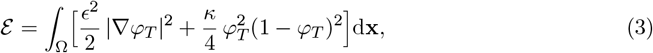

where *κ* is the diffuse interface width and *ϵ* the adhesion energy scale [28]. The chemical potential *ν* = *δE/δφ*_*T*_ follows from the variational derivative of (3):

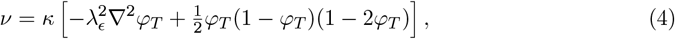

which drives the diffusive flux ***J***_*i*_ = *m*_*i*_*φ*_*i*_ ∇*ν*, with *m*_*i*_ as the cell motility.

Advection is driven by a Darcy-like solid velocity **u**_*s*_ that couples growth to mechanical relaxation [47]:

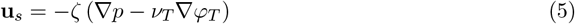

where *p* is an effective mechanical pressure. Volume conservation ∇· **u**_*s*_ = *S*_*T*_ gives a Poisson equation for *p*:

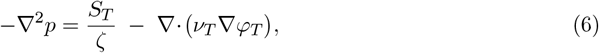

subject to homogeneous Neumann boundary conditions. The net source *S*_*T*_ = *S*_*V*_ + *S*_*N*_ is specified below.

We introduce a general relative nutrient concentration field 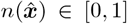. We assume this field equilibrates rapidly relative to growth timescales [28, 47], justifying a quasi-steady reaction–diffusion equation with density- and uptake-dependent diffusivity hindrance:

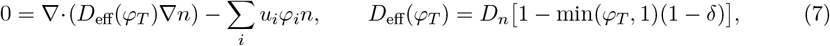

where *u*_*i*_ is the uptake rate of population *i, D*_*n*_ the baseline diffusivity, and *δ* ∈ [0, 1] the minimum diffusion fraction. In the present viable–necrotic model, uptake is provided by the viable population, so the sum reduces to *u*_*V*_ *φ*_*V*_ *n*. Diffusion is unimpeded (*D*_eff_ = *D*_*n*_) outside the spheroid and reduced to *δD*_*n*_ in dense tissue [49, 50]. The nondimensionalisation of the model in Supplementary Information shows how this equation is reduced to a characteristic exponential penetration-depth scale, and that the effect of *δ* cannot distinguish between density hindrance and an increase in the consumption rate. As such, it is interpreted as a parameter that jointly represents the total effect of the tissue (mechanical or biological) on nutrient diffusion.

### 2.2 Source and sink terms

All cell fate pathways are deterministically encoded in *S*_*i*_. We decompose each source term into incoming (proliferation) and outgoing (death) contributions: *S*_*i*_ = *I*_*i*_ − *O*_*i*_ (the generalised population-transfer structure is provided in Supplementary Information). We introduce two modifications to the standard form used in Wise *et al*. [47]: a global necrotic feedback term and a continuous nutrient-dependent death switch.

#### Proliferation

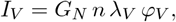

where *λ*_*V*_ is the effective proliferation rate and *G*_*N*_ is a global necrosis-mediated inhibition factor.

#### Cell death

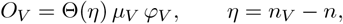

where *µ*_*V*_ is the effective viable cell death rate, and Θ(*η*) is the smooth nutrient switch:

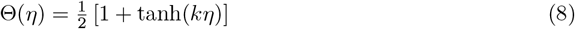

where *k* sets the switch steepness. This replaces the discontinuous Heaviside function commonly used in compartment models [33, 39] with a smooth transition consistent with the continuum framework. The value of *k* directly controls the width of the viable-to-necrotic transition zone: large *k* produces a sharp boundary resembling the compartment idealisation, while small *k* yields a broad gradient. Since Θ couples to the local nutrient field *n*(**x**), which itself depends on the spatial distribution of cells through consumption and hindrance, the switch steepness has a spatially dependent effect on internal structure that cannot be captured by a fixed compartment width. This provides an additional degree of freedom for shaping the inhibited region and encoding sensitivity to nutrient gradients.

#### Global necrotic feedback

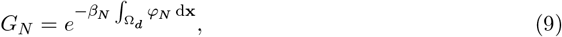

where Ω_*d*_ denotes the spatial domain in *d* dimensions and *β*_*N*_ controls feedback strength. This captures an aggregate inhibitory effect of accumulated necrotic material on further expansion [51, 33, 39].

#### V–N source terms

Restricting to viable and necrotic cell types, the source terms reduce to:

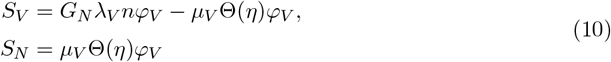

The vectorised form of these equations (Supplementary Information) enables straightforward extension to additional cell types, such as quiescent type or stem cells.

#### Drugged Conditions

Within this structure, drugged conditions can be modelled trivially. Assuming the existence of a relative drug concentration field *d*, mirroring Equation 7, we define an *activity function a*_*i*_(*d*) which describes the strength of the interaction between the drug field and cell population *i*. Within the existing V–N source term structure, we perturb the key mechanistic biological parameters, with a proportionality to the activity function.

- Growth Suppression: 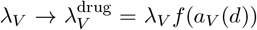 where *f* is a monotonically decreasing function of the activity.
- Direct Death: 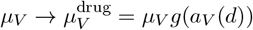 where *g* is a monotonically increasing function of the activity.
- Nutrient Sensitisation: 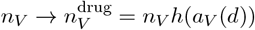 where *h* is a monotonically increasing function of the activity.

Full prescription of these drugged source terms is provided in Supplementary Information.

### 2.3 Non-dimensionalisation

Spatial coordinates are non-dimensionalised by the dataset-specific initial spheroid radius *R*_0_, so that 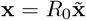. The nutrient transport scale is described by an effective penetration depth, motivated by the steady-state reaction–diffusion balance. The radius-scaled nutrient equation contains the length ratio 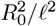. We then rewrite this ratio using a dimensionless sweep parameter:

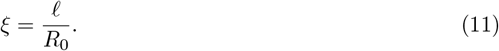

The simulation library is generated in non-dimensional form and indexed by *ξ*. Doing this allows a single simulation library to be mapped to any experimental condition through appropriate spatial scaling, since *ξ* captures the relative balance of nutrient diffusion and consumption that shapes internal structure. The entire non-dimensional model is presented in Supplementary Information. Nominal values of *ξ* describe the relative penetration depth of the nutrient with respect to the initial radius. When *ξ* ≥ 1, this describes a regime in which the spheroid radius is less than the characteristic penetration depth of the extracellular nutrient, implying near-saturation of nutrient throughout the spheroid. For *ξ* ≤ 1, this describes the regime where the characteristic penetration depth is smaller than the initial radius. This is the expected regime, where nutrient depletion towards the centre of the spheroid (*r < R*_0_ − *ℓ*) induces nutrient-limited structures.

### 2.4 Model and Numerical Implementation

The experimental data used for the target calibration radii were measured from two-dimensional experimental image planes of the equatorial section of the spheroid [37], therefore solve the continuum fields on a two-dimensional Cartesian domain and compare the resulting image-plane radii directly with those measurements.

For clarity, the simulation can be summarised as a sequence of field evaluations and updates.

At each timestep, the system is solved in the following order:

- Necrotic feedback, Eq. (9).
- Nutrient switch and source terms, Eqs. (8)–(10).
- Chemical potential, Eq. (4).
- Pressure equation, Eq. (6).
- Solid velocity, Eq. (5).
- Viable and necrotic population update, Eq. (2).
- Quasi-steady nutrient update, Eq. (7).

The entire model and the calibration infrastructure was implemented using Python. The equations are solved using a uniform Cartesian grid with second-order finite-difference operators, implemented in NumPy with Numba acceleration. Time integration uses a Crank-Nicolson scheme where the stiff fourth-order term is handled implicitly via a discrete cosine transform (DCT) [52], while all other terms remain explicit. Equation 6 is solved using the same DCT approach. Equation 7 is solved via fixed-point iteration using the SciPy conjugate-gradient solver. All inner loops are compiled with Numba jit for single-node parallel execution. On a single Apple Silicon core (M4 Pro), a 100^2^ simulation with 40 time steps completes in approximately 0.3 seconds. The reported sweep throughput is approximately 850,000 simulations every 24 hours with approximately 10 parallel workers. The nondimensionalisation described in Section 2.3 enables a single simulation library indexed over *ξ*_*i*_ to be mapped to any experimental condition through appropriate spatial scaling. A full description is provided in Supplementary Information.

Radial observables are implicitly computed from the simulated fields, using a contour-based method averaging the distance from the total tumour (*φ*_*T*_) centre of mass to boundary points where the target field exceeds a specified threshold.

## 3 Methodology – Model Calibration to Experimental Data

With the mathematical and numerical framework established, the next objective is to calibrate the model against experimental measurements of tumour spheroids. The core aim is to reproduce classical untreated tumour spheroid growth dynamics [33]. Because the continuum model does not impose explicit compartment boundaries, we treat the mapping of the inhibited radius as an exploratory model-to-data comparison: we calibrate the model against the total and necrotic radii, then evaluate multiple candidate definitions for the intermediate region against the experimentally measured trajectory. This comparison identifies which continuum observable gives the strongest relative agreement under the CTM; it is not intended to establish the biological identity of the experimental compartment.

The full inverse model calibration contains *seven* parameters governing biological and nutrient dynamics, together with the library sweep coordinate *ξ*. Preliminary exploratory simulations identified the parameters that have negligible influence on the macroscopic radii within these experimental time frames and were therefore fixed to biologically plausible values (reported in Supplementary Information). The remaining parameters were found to have measurable effects on the total, inhibited, and necrotic radii in preliminary sweeps and were treated as free: ***θ***_*b*_ = {*λ*_*V*_ , *µ*_*V*_ , *n*_*V*_ , *δ, ζ, β*_*N*_ , *k*}. The coordinate *ξ* is swept alongside these parameters so that one universal simulation library covers the six dataset-specific initial-radius regimes; it indexes the represented physical scale rather than introducing an additional biological mechanism. The prior distributions and nominal values for the biological and nutrient parameters were informed by preliminary sweeps, literature values, and biological plausibility, and are reported in Supplementary Information.

### 3.1 Experimental data

Calibration was performed using longitudinal confocal imaging data from the WM983b and WM793b melanoma spheroid lines at 10000, 5000, and 2500-cell seeding densities, as reported by Browning et al. [37]. Their analysis pipeline combines bright-field segmentation with FUCCI-based fluorescence to quantify: (i) the total spheroid radius (*R*_*T*_), (ii) the inhibited-region boundary (*R*_*I*_), and (iii) the necrotic-core radius (*R*_*N*_) [37]. Browning *et al*. define the inhibited boundary using a 20% maximum mAG fluorescence threshold, and the necrotic boundary via a texture-based classifier. Browning’s inhibited radius is an experimentally operational definition; it is not uniquely tied to a single state variable of the CTM, motivating the observable-selection approach used here.

### 3.2 Objective function and likelihood

To compare the simulated radius trajectories to the experimental data, we adopt an uncertainty-weighted root-mean-square error (WRMSE):

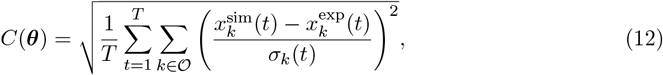

where *O* = {*R*_T_, *R*_I_, *R*_N_} denotes the total, inhibited, and necrotic radii, *x*_*k*_(*t*) denote the experimental and simulated data points at time *t* for observable *k*, and *σ*_*k*_(*t*) is the experimental standard error. Under independent Gaussian error assumptions, the log-likelihood is a monotone decreasing function of *C*^2^(***θ***) [53], so minimising *C*(***θ***) is equivalent to maximising log ℒ (***θ***). The later extension of Browning’s model to a time-evolving Greenspan-type ODE model [39] was reimplemented and evaluated on identical datasets for direct model performance comparison.

### 3.3 Calibration pipeline

Model calibration proceeded through three sequentially informed stages, each addressing a specific challenge of the inverse problem: a computationally expensive forward model, the absence of gradient information, and structural uncertainty in the definition of the inhibited radius. All parameter prior distributions for all calibration experiments are reported in Supplementary Information.

#### Stage 1 — Global exploration via Latin hypercube sampling

We first built a coarse but space-filling map of the parameter–observable relationship using Latin hypercube sampling, which partitions each parameter prior into equiprobable intervals and draws one sample per interval to ensure uniform coverage. Each sampled parameter vector was passed through the simulation engine and scored using the objective (Equation. (12)).

Because the CTM has no explicit inhibited compartment, multiple candidate definitions exist (necrotic-to-total fraction thresholds, nutrient concentration isosurfaces, local proliferation and death strength isosurfaces). At this stage, all candidates were computed and scored independently to compare model-internal mappings and identify plausible parameter regions, without committing to a single definition. Candidate thresholds were evaluated every 5% within the necrotic-density family, nutrient-concentration family, and relative positive-source, negative-source, and pressure families. These global comparisons were used to initialise the subsequent exploratory Bayesian analysis; candidate weights were not interpreted as biological probabilities.

#### Stage 2 — Particle SMC for posterior sampling

Regions of parameter space from Stage 1 that provided strong fits were used to reduce the breadth of the parameter priors defined for a sequential Monte Carlo (SMC) sampler, based on the algorithm developed by Jeremiah *et al*. [54]. At each round within the prior distributions, a population of particles (points in parameter space) were assigned importance weights proportional to their posterior density, resampled when the effective sample size fell below a threshold, and moved via random-walk Metropolis–Hastings steps.

Exploratory Bayesian SMC runs initially included the full set of inhibited-radius candidates under equal prior weights. The inhibited-radius definition was treated as a latent discrete parameter and marginalised in the likelihood, allowing the SMC posterior to weight each candidate according to its consistency with the data. Because different candidate definitions produced distinct regions of parameter space, these all-candidate runs were used to compare candidate-associated basins rather than to define the final calibration directly.

The most promising candidate-associated basin for each dataset was then selected from the exploratory SMC results. A subsequent focused SMC run fixed the selected inhibited-radius candidate and refined its parameter basin using narrower bounds. We summarised the resulting posterior spread using multiples of the posterior standard deviations and refer to these summaries as posterior plausible intervals rather than formal Bayesian credible intervals. These plausible intervals were used to define local parameter ranges and inform the bounds of the third stage of calibration: profile likelihood analysis. All resulting candidate support and parameter summaries should therefore be interpreted as conditional on the CTM and on this exploratory basin-selection workflow.

#### Stage 3 — Profile likelihood analysis

To complement the SMC posterior summaries, we computed profile likelihoods for each parameter of the identified best fits. For parameter *θ*_*k*_, the profile log-likelihood is 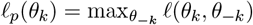, obtained by fixing *θ*_*k*_ at a grid of values and re-optimising over all remaining parameters. Likelihood-based confidence intervals were extracted using Wilks’ theorem: 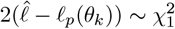, where 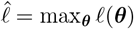, with the Δ*ℓ* = 1.92 threshold corresponding to approximately 95% confidence. These profile likelihood sweeps were used to finalise the best-fit points and categorise parameter identifiability. The profile-likelihood confidence intervals were then compared with the SMC posterior plausible intervals to check that both summaries referred to the same local basin.

This three-stage programme produces a combined view of the calibrated model: Stage 1 provides the global cost landscape allowing for identification of local minima; Stage 2 yields posterior samples, MAP estimates, posterior means, posterior standard deviations, and the marginalised distribution over inhibited-radius definitions; Stage 3 provides a complementary local likelihood-based identifiability assessment by comparing profile-likelihood confidence intervals with posterior plausible intervals.

## 4 Results

The calibration pipeline described in Section 3 was applied to the Browning *et al*. spheroid datasets. The final accepted fits combine SMC basin-finding sweeps with profile-likelihood confirmation. The SMC sweeps were used to identify plausible fit basins, posterior structure, and candidate intermediate-radius definitions. Profile likelihoods were then used to obtain local confidence intervals and assess practical identifiability within the selected basins. The reportable parameter values for all six datasets are summarised in Supplementary Information.

### 4.1 Latin Hypercube Exploration of the Parameter Space

Global exploration was performed by generating approximately 500,000 parameter combinations per *ξ* value via Latin hypercube sampling, spanning the seven biological/mechanical parameters and the library scale coordinate. Each parameter vector was passed through the simulation engine and scored via the WRMSE (Eq. 12). Figure 2 summarises the resulting objective landscape, while Figure 3 visualises the competing field-derived intermediate-radius candidates and their representative exploratory comparison.

The objective landscape revealed initial parameter fits to the experimental data, as well as insight into the structure of the model. In Figure 2(B), the parameter-pair landscapes show this structure, with *λ*_*V*_ and *β*_*N*_ showing signs of positive correlation. *δ* and *n*_*V*_ , and *n*_*V*_ and *k*, show inverse correlative structures through intersecting objective valleys, and *µ*_*V*_ vs. *β*_*N*_ shows an example of two parameters that are not individually resolved from one another. *β*_*N*_ is constrained to a small partition of the x-axis, but many values of *µ*_*V*_ produce similar resulting experimental fits. *n*_*V*_ and *λ*_*V*_ , and *λ*_*V*_ and *k*, are examples of well-resolved minima, where a relatively small parameter region produces the strongest fits to the experimental data.

This global exploration and comparison to the experimental data was handled by the two-layer database structure described in Figure 2(D). This handled comparisons to all experimental datasets at simulation time, across all simulation observables. This allowed us to extract initial estimates of parameter combinations for each dataset, which ranged from approximately 0.8–1.7 on the aggregate standardised score defined in Eq. 12. These initial estimates span a range of observables attributed to the inhibited radius, including the interior necrotic structure (continuum density profile), relative positive source intensity (where proliferation is occurring), relative negative source intensity (where cell death is occurring), nutrient concentration, and relative pressure strength across all six datasets. Figure 3(A) shows how these candidate families correspond to distinct radial structures in a single simulated spheroid.

Notably, the different inhibited radius candidates for the same dataset often sit within different regions of parameter space, indicating different model modalities under the objective function across different inhibited-radius definitions. We therefore ran exploratory Bayesian SMC sweeps over the full inhibited-radius candidate set. Candidate thresholds were evaluated every 5% and assigned equal working weights, allowing the SMC procedure to compare the competing definitions through their likelihoods while exploring their associated parameter basins. Figure 3(B) gives a representative WM983b 5k example: relative support is concentrated almost entirely in the necrotic-structure family, and within that family the 70% threshold is strongly preferred. Across the datasets, the exploratory SMC runs consistently identified promising basins associated with the necrotic-structure observables, with some relative support also assigned to positive- and negative-source structures. The pressure structures appeared as an outlier in one dataset, where the experimental inhibited-radius measurements contain greater variability. The most promising candidate-associated basin for each dataset was then selected for focused SMC refinement. These comparisons are conditional on the CTM and are not interpreted as biological probabilities.

### 4.2 SMC and Profile-Likelihood Fits Across Datasets

For each dataset, an exploratory Bayesian SMC run first weighted the full inhibited-radius candidate set while sampling the associated parameter space. The most promising candidate-associated basin was then selected, and a subsequent focused SMC run refined that basin with the chosen candidate fixed. The selected candidates varied slightly across datasets but were consistently aligned with mid-to high-density necrotic regions (*φ*_*N*_ ≈ [0.5, 0.7]). Across all six datasets, the calibrated simulations achieved aggregate WRMSE values between 0.489 and 1.236 on the standardised score defined in Eq. (12). The strongest fits were identified from the WM983b datasets, with an average aggregate WRMSE of 0.634, whereas the WM793b datasets achieved an average aggregate WRMSE of 1.010. To assess local parameter resolution within the accepted basins, we computed profile likelihoods for each parameter, following established profile-likelihood approaches to practical identifiability and finite-sample interval estimation [53, 55, 56]. For each parameter *θ*_*k*_, a fixed grid was evaluated with re-optimisation over the remaining parameters at each grid point. Confidence intervals were then read using the Δ log ℒ = 1.92 threshold, corresponding to an approximate 95% interval.

Figure 4 presents the results of the WM983b 5k dataset as the representative model structure and strongest overall calibration. Across multiple SMC parameter sweeps, independent seeds converged to the same basin, and profile likelihoods were computed for all biological and nutrient-associated parameters. The accepted fit uses a necrotic-structure inhibited-radius proxy at the 70% threshold. The posteriors and basin plots (Figure 4(A,B)) offer insights into the compensatory structure identified in the model. Across all six datasets, a compensatory structure existed between the nutrient transport and death mechanisms. Consistent correlations between *k, n*_*V*_ , *δ*, and *µ*_*V*_ indicate that these parameters were not all individually resolved, but the parameter combinations were constrained. This appears in the posteriors as wider distributions, where these parameters occupy broader regions of parameter space while other parameters compensate. We also see some correlation between the effective growth rate *λ*_*V*_ and growth inhibition *β*_*N*_ , with both parameters being part of the same source term and having opposing effects in Eq. 10, such that an increase in both parameters keeps the source term behaviour similar. The pressure constant was not resolved across the datasets. Being the only purely physical constant swept over, there was likely no mechanism within the experimental design that could resolve this parameter. An experiment that investigates how spheroids grow in different external conditions (free media, hydrogels, scaffolds, etc.) would likely provide the type of variation needed to resolve a parameter with the pressure constant’s role in the model.

Across all six datasets, all parameters except *ζ* met one of the manuscript’s local resolution criteria within the accepted calibration basins: parameters were classified as *strongly resolved* when the profile-likelihood confidence-interval width was < 25% of the MAP, or *resolved* when it was < 50% of the MAP. These labels describe the width of the profile-likelihood interval relative to the fitted MAP under the stated definitions. The use of relative interval width is consistent with previous practical-identifiability analyses that compare parameter resolution using finite confidence intervals or relative confidence-interval sizes [53, 57, 55]; the 25% and 50% cut-offs are operational criteria defined for this study, not universal identifiability thresholds. Accordingly, these labels are not claims of independent or global identifiability. The classifications are summarised by the resolution, posterior, and profile-likelihood panels in Figure 5(A,D,E). Several intervals should also be interpreted with care because some parameters are constrained by one-sided physical bounds or saturation behaviour rather than symmetric interior likelihood curvature, and correlated parameter combinations may remain compensatory.

Figure 4(C) presents this fit to the experimental data with corresponding experimental uncertainty bands. The fit to the data is very strong (*C* = 0.492 on the aggregate standardised score) across all three radii, with all experimental data points sitting directly on the simulated trajectory or within the 68% experimental uncertainty band.

### 4.3 Comparison with the Greenspan-type Model

To contextualise the performance of the continuum model, we re-implemented the Greenspan-type ODE model of Murphy *et al*. [39] and calibrated it under identical fitting criteria. The WRMSE objective and the Gaussian log-likelihood are equivalent for model ranking under the shared error model, so both models were compared using the same data and uncertainty weighting. The continuum model is nevertheless more complex because it contains additional field dynamics and an exploratory model-internal mapping for the inhibited radius. The Greenspan model, which explicitly parameterises the three canonical compartments, achieved an average aggregate WRMSE of *C*_G_ = 1.35, with a global best fit of 0.837. The continuum model achieved a database-wide average aggregate WRMSE of 0.746, and a global best fit of 0.505, with an average dataset-wise improvement of approximately 40%; the matched trajectories are shown in Figure 5(C). The reportable profile-supported rows in Supplementary Information are a defined subset of the accepted fits and are therefore not expected to reproduce this database-wide average exactly.

This comparison is important because the continuum model in its formulation does not identify the existence of the classical spheroid compartments. Instead, the relevant radii are extracted from simulated fields and selected through the calibration objective. The stronger result above is not just a strong fit to the experimental data: the classical three-phase spheroid growth structure can be recovered within the continuum model, while fitting the data better than a model constructed around that structure.

Two particular parameters required additional consideration for generating profile likelihoods. In particular, *k* acts as a saturation parameter, where *k* → ∞ ⇒ Θ → ℋ, and ℋ is the Heaviside step function. This implies that sufficiently large values of *k* can produce similar sharp-switch behaviour in the objective function. Furthermore, for the WM983b 2.5k, WM793b 2.5k, and WM793b 5k datasets, *β*_*N*_ attained its optimum at or near the lower physical boundary *β*_*N*_ = 0, as reflected in the corresponding profile-likelihood ridges in Figure 5(E). Because these optima occur at physical parameter boundaries, the regularity assumptions behind a symmetric Wilks-based confidence interval are not satisfied. We therefore report one-sided profile-likelihood intervals for these boundary-constrained parameters. Complete parameter values and confidence interval ranges are reported in Supplementary Information.

The strongest cross-dataset pattern is that WM983b and WM793b do not use the necrotic/inhibited-radius structure in the same way. WM983b 5k and WM983b 10k both support nonzero *β*_*N*_ , while WM983b 2.5k fits well with *β*_*N*_ close to zero. By contrast, WM793b 2.5k and WM793b 5k both support low or near-zero *β*_*N*_ , and WM793b 10k only supports nonzero *β*_*N*_ in a harder, more compensatory basin. This cross-dataset structure is visible in the posterior and profile-likelihood summaries in Figure 5(D,E). It indicates that the impact of necrotic feedback is much stronger in the WM983b line than in the WM793b line, where it appears to become nonzero only at the higher seeding density, where necrosis is more prominent.

The nutrient density-hindrance parameter *δ* also varied across datasets. Assuming consistent tissue density hindrance across datasets, a low value for *δ* indicates a higher rate of nutrient consumption and higher values indicate less consumption. WM983b 2.5k and WM793b 5k supported very low hindrance, WM983b 10k and WM793b 2.5k supported low but nonzero hindrance, WM983b 5k supported moderate hindrance, and the accepted WM793b 10k basin supported higher hindrance. This pattern, together with its correlation structure, is shown across the posterior and correlation panels in Figure 5(B,D). It is not fully consistent across cell lines or seeding densities, but higher seeding densities appear to support higher values of *δ*, indicating less nutrient consumption. This aligns with the idea that a larger initial radius, where tissue penetration has a stronger impact on nutrient transport, would affect the cells’ consumption rate. Nutrient-response parameters were not independent, so variations in these nutrient-associated parameters need to be interpreted through their relationships with correlated parameters.

There is also a distinction in fit quality between datasets. The average MAP aggregate WRMSE for the WM983b datasets is 0.633, compared to 1.001 for the WM793b datasets; the dataset-specific fits are shown in Figure 5(C). One plausible explanation is a mismatch between the data and the initial conditions of the simulation configuration. The simulation configuration assumed canonical spheroid growth and was therefore seeded with purely viable cells, instantiating an effective replicate of phase 1 (Figure 1). The WM793b datasets consistently measured a nonzero inhibited radius at the initial data point, potentially due to external influence or cell sensitivity, which would reduce fit quality under a purely viable initial condition. After initial calibration sweeps in the assumed canonical regime, we identified the best performing inhibited radius candidate, and performed a sweep across different initial inhibited radii conditions. We kept all other calibrated parameters fixed and reoptimised over the initial condition configurations (Figure 4 A).

**Figure 1:**
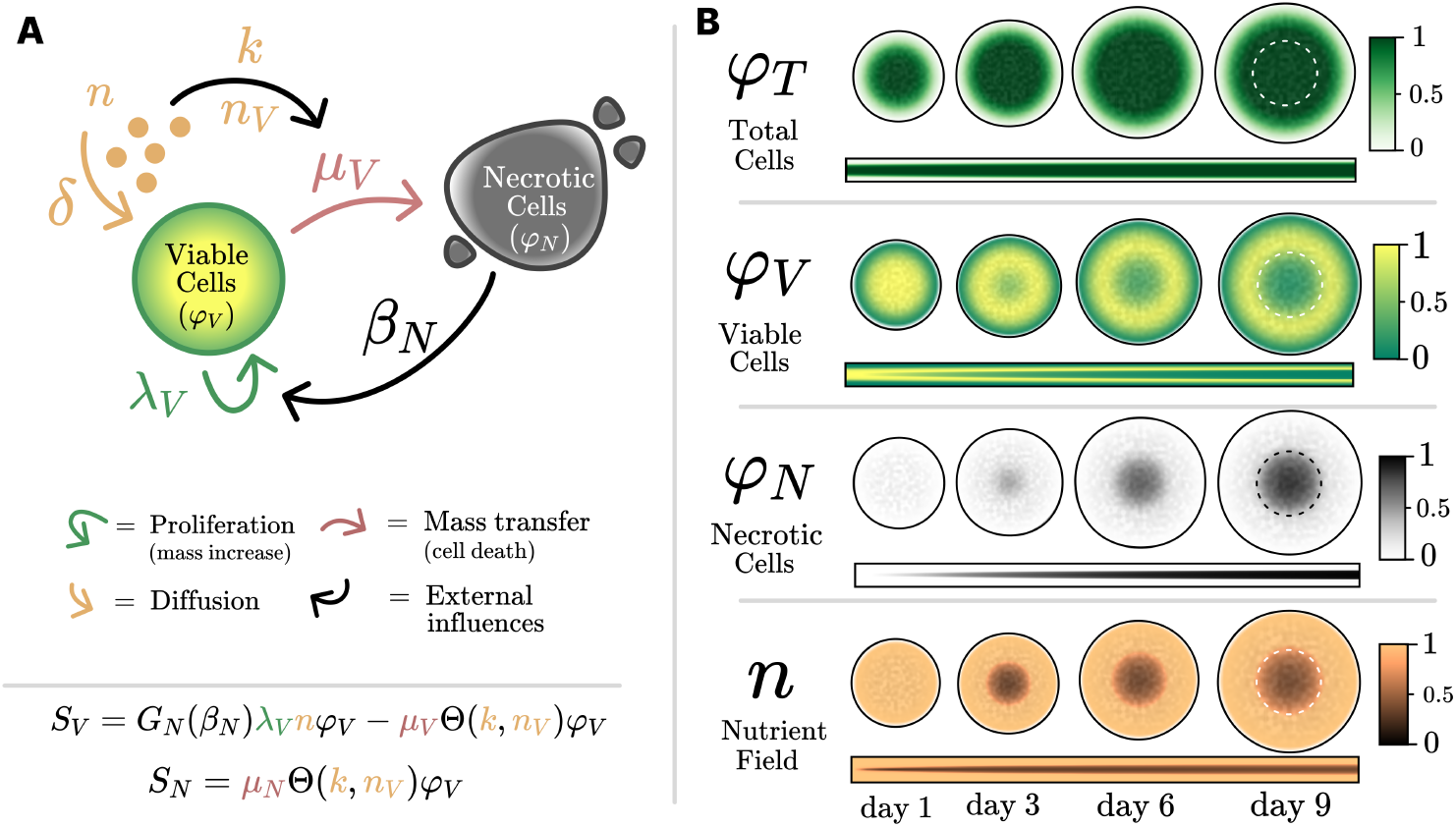
Cell Schematic and Emergent Spheroid Growth Structure. (A) Demonstrates the cell pathway dynamics, and the model parameters impact on those pathways. (B) Model implementation demonstrating the evolution of the cell and nutrient fields and the emergence of the characteristic three phase growth structure. Day 1 demonstrates an entirely viable seeded spheroid (phase 1), day 3-6 demonstrates the development of the internal structure due to nutrient depletion (phase 2), and day 9 demonstrates the emergence of a necrotic core due to sustained nutrient deprivation (phase 3).

### 4.4 Drug Response

The strongest motivation for developing this model is to build a framework that allows the internal structure of spheroids to be studied under drug-treated conditions. Classical Greenspan-type models, like the comparison model [39], cannot support some of the expected structures that would form under drug-treated conditions without major reformulation. Structurally, these models support an ordered, three-layer structure consisting of a viable rim, necrotic core, and intermediate region, limiting their flexibility to represent different structures where necrotic regions may form near the exterior. Here we introduce a continuum drug field into the existing model as a proof-of-concept demonstration of the model’s extensibility rather than as a validated drug-response inference framework.

We implement a second quasi-steady reaction–diffusion equation in the same form as Eq. 7, with approximately half the nutrient diffusion constant to simulate larger drug molecules. This drug field *d* is coupled to the existing source terms in several ways. We implement five source-term variants representing three impacts of the drug field on viable cells. We consider direct death (dd), growth-rate suppression (gs), and nutrient sensitisation (ns) through simple scalings of the associated parameters (*λ*_*V*_ , *µ*_*V*_ , and *n*_*V*_). We then consider direct death combined with growth suppression (dg), followed by all impacts combined (c). We present the impact at three relative strengths and three diffusion rates to simulate varying response magnitudes and penetration depths. All drug-effect simulations are performed around a single calibrated MAP parameter set, so posterior uncertainty is not propagated into the drug-response figure. We chose the WM983b 2.5k seeding density so that the width of the viable region with respect to the necrotic core was the smallest, maximising the resolution of the viable cell impact. We apply a delayed instantiation of the drug field to allow the spheroid to mature and develop internal structure before drug application. The results of the drug field experiments are displayed in Figure 6. The full drug-response implementation is detailed in Supplementary Information.

The drug-attributed viable cell loss displayed in Figure 6(A) represents the magnitude of viable cell loss in each region that can be attributed to the drug field. Each condition shows a distinct profile across the radial coordinate of the spheroid. Direct death appears to affect the spheroid earlier and more strongly, with a deeper impact relative to the other conditions. Growth suppression produces a more delayed response, with its strongest magnitude occurring near the end of the assay, and does not penetrate as deeply because of the slowing of radial expansion. Nutrient sensitisation has the highest magnitude response, and its impact lasts longer than direct death. Nutrient sensitisation penetrates deeper into the spheroid than growth suppression, but not as deeply as direct death. The combination treatments also have distinct profiles. Direct death combined with growth suppression combines the early, deeper response from direct death with the delayed impact from growth suppression, making the effect remain stronger for longer than in the purely direct-death condition. The all-effects combination further follows this trend with a longer, deeper profile and a wider, stronger overall impact. The total time-integrated viable cell loss attributed to the drug effect shows that the combination effects produce the most death, followed by direct death, nutrient sensitisation, and finally growth suppression.

The analysis of drug response within this model would likely rely on these viable-cell-loss impact heatmaps. Here, drug-attributed viable cell loss is defined as the simulated reduction in viable-cell source contribution relative to the untreated calibrated baseline under the same model state. The distinct shape and magnitude of these profiles provide insight into how viable cells in the system were affected by the imposed drug field. Experimentally, this may provide the ability to quantitatively profile the magnitude of drug impact (time-integrated drug-attributed viable cell loss), the magnitude of drug penetration, and, importantly, the mechanism from which the viable cell loss was sourced.

Figure 6(B) demonstrates the effect of the drug through the classical spheroid structural lens. We use the calibrated definition for the inhibited radius of the WM983b 2.5k dataset and track how it, the necrotic radius, and the total radius vary under different drug mechanisms. All drug-treated conditions reduce the total radius, with the combined and direct-death mechanisms producing the greatest perturbation. The effect on the inhibited radius appears more nonlinear, with some conditions causing spikes and later reductions. The necrotic radius generally increases under drug-treated conditions, with a local peak followed by a reduction relative to the control. This can be attributed to an initial drug-driven increase, followed by the control necrotic radius overtaking the drug-treated conditions because of the drug’s global effect on spheroid growth.

Figure 6(C) is a radial profile of the viable cell population, using a diverging colourmap to implicitly visualise the calibrated inhibited radius. The transition between red and blue in the control is exactly the inhibited radius for the WM983b 2.5k dataset, and how this region varies therefore provides insight into the breaking of the classical layered structure. Under some stronger drug-treated conditions, such as direct death and the combination mechanisms, this transition “radius” is altered and widened by the impact of viable cells dying due to the drug. This displays a fundamental variation from the layered model structure under drug-treated conditions: one-dimensional scalar radii under untreated conditions transform into intermediate regions. As with the drug-attributed loss profiles, categorising the viable-cell profile itself under drug-treated conditions will give further insight into the impact and effect of the applied drug.

## 5 Discussion

The central result of this work is that a general continuum cell-growth model can be calibrated to classically described spheroid growth while locally constraining biological and nutrient-related parameter combinations, as well as comparing macroscopic field observables as model-internal proxies for the experimentally derived “inhibited radius”. Without prescribing compartments in the model structure, the continuum model recapitulated the canonical three-phase growth described by Greenspan [33] (Figure 1) and improved average fit quality by approximately 40% compared to a Greenspan-type model [39], which models the compartmental structure directly (Figure 5(A)). Exploratory Bayesian SMC runs compared the competing inhibited-radius candidates and identified promising basins associated with the necrotic structural family, as illustrated by the candidate fields and relative support in Figure 3; the selected basins were then refined in focused SMC runs. This comparison is conditional on the CTM and does not claim to identify the biological nature of Browning’s experimental boundary.

Phase-field and diffuse-interface tumour models have been developed and refined over nearly two decades, yet their application has remained generally theoretical: demonstrating emergent morphological features, exploring parameter regimes qualitatively, or benchmarking numerical methods against synthetic targets [47, 48, 29]. Direct, quantitative calibration of classical spheroid structure against experimental growth data has been notably rare within the literature. Lima *et al*. [58] demonstrated that a continuum formulation could be calibrated to time-resolved tumour measurements, although without spatial calibration and within the context of two-dimensional cell culture systems rather than three-dimensional multicellular spheroids. To our knowledge, no phase-field model has been systematically calibrated against longitudinal, multi-observable spheroid imaging data of the kind reported by Browning *et al*. [37]. Several factors contribute to this gap. The experimental datasets needed to constrain such models (longitudinal, multi-observable measurements that resolve not only total size but also internal compartmental structure) have only recently become available through advances in high-content imaging and fluorescence-based segmentation [37, 45, 18]. Communities developing phase-field theory and those generating spheroid data have appeared to operate largely in parallel, with one-dimensional radially symmetric models, originating from the Greenspan-type ODE approaches, serving as the primary modelling tools within the experimental spheroid literature [33, 34, 37, 39]. The present work bridges this gap by combining a computationally tractable general cell-growth continuum model under spheroid-style initial conditions with a multi-observable calibration protocol applied to the same experimental datasets used to validate established spheroid models (Figures 2, 3, and 5), thereby providing a direct and controlled comparison between frameworks.

**Figure 2:**
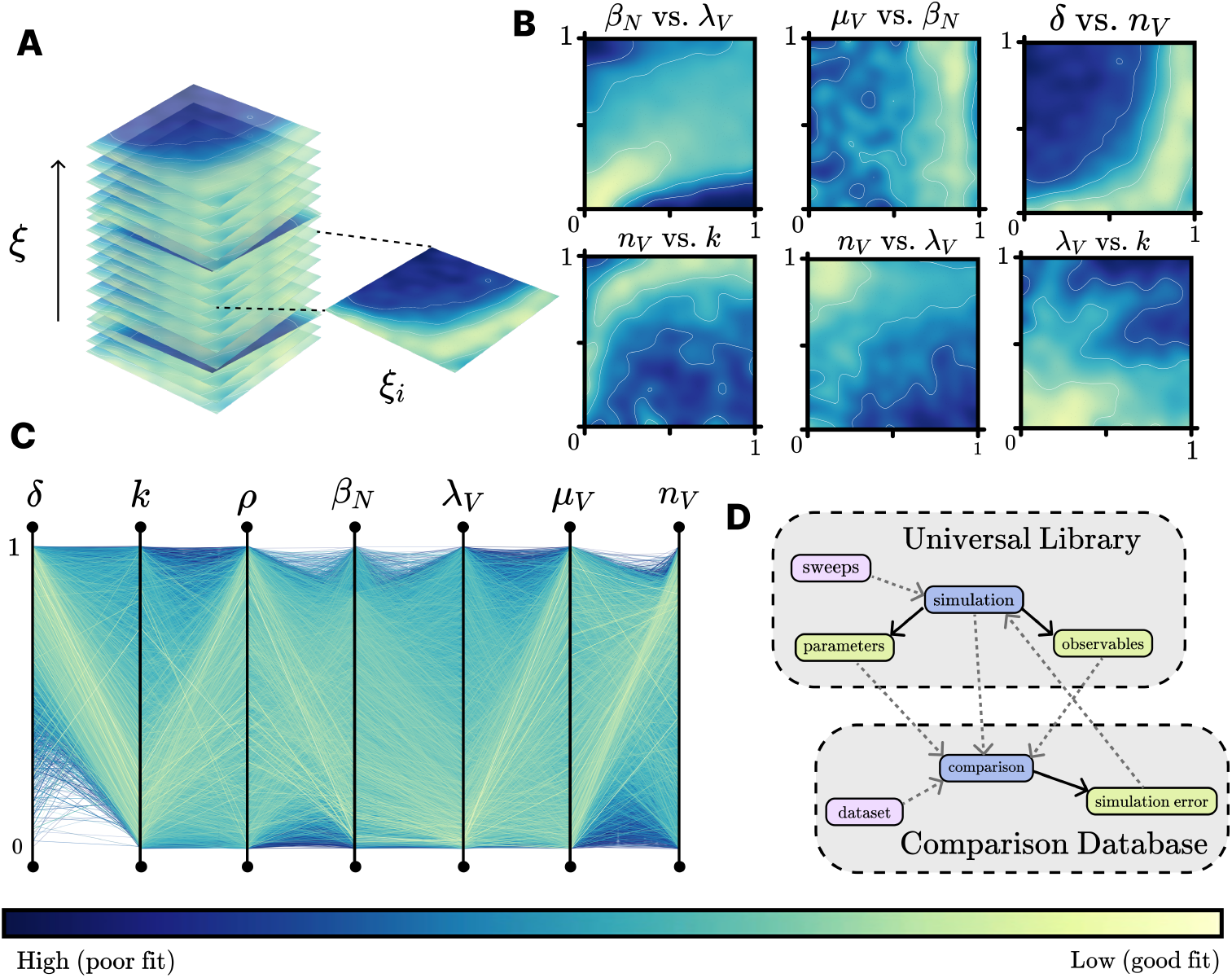
Global exploration via Latin hypercube sampling for the WM983b 5k dataset. (A) Depicts a representation of the universal database from which a “slice” is removed for the corresponding dataset’s *ξ* value (relative nutrient penetration depth). (B) Two-dimensional parameter-pair mixed observable objective landscapes (presents WRMSE for all inhibited radius candidates, and parameter label corresponds to y-x, bounds can be found in Supplementary Information). Light regions correspond to areas of the parameter space which produce strong fits, while dark regions represent weak parameter combinations. (C) Parallel coordinate plot of parameter values, revealing constrained parameters as tightly coupled strands and unconstrained parameters as diffuse distributions. (D) Depicts the process of database storage for the universal databases. Sweeps are run, producing simulations, with a parameter and observable set. These simulations are then compared to specific datasets, in a supplementary “comparisons” database. The compared simulations have an associated simulation ID such that WRMSE values can be attributed to a particular simulation in the universal database, and as such a parameter and observable set.

**Figure 3:**
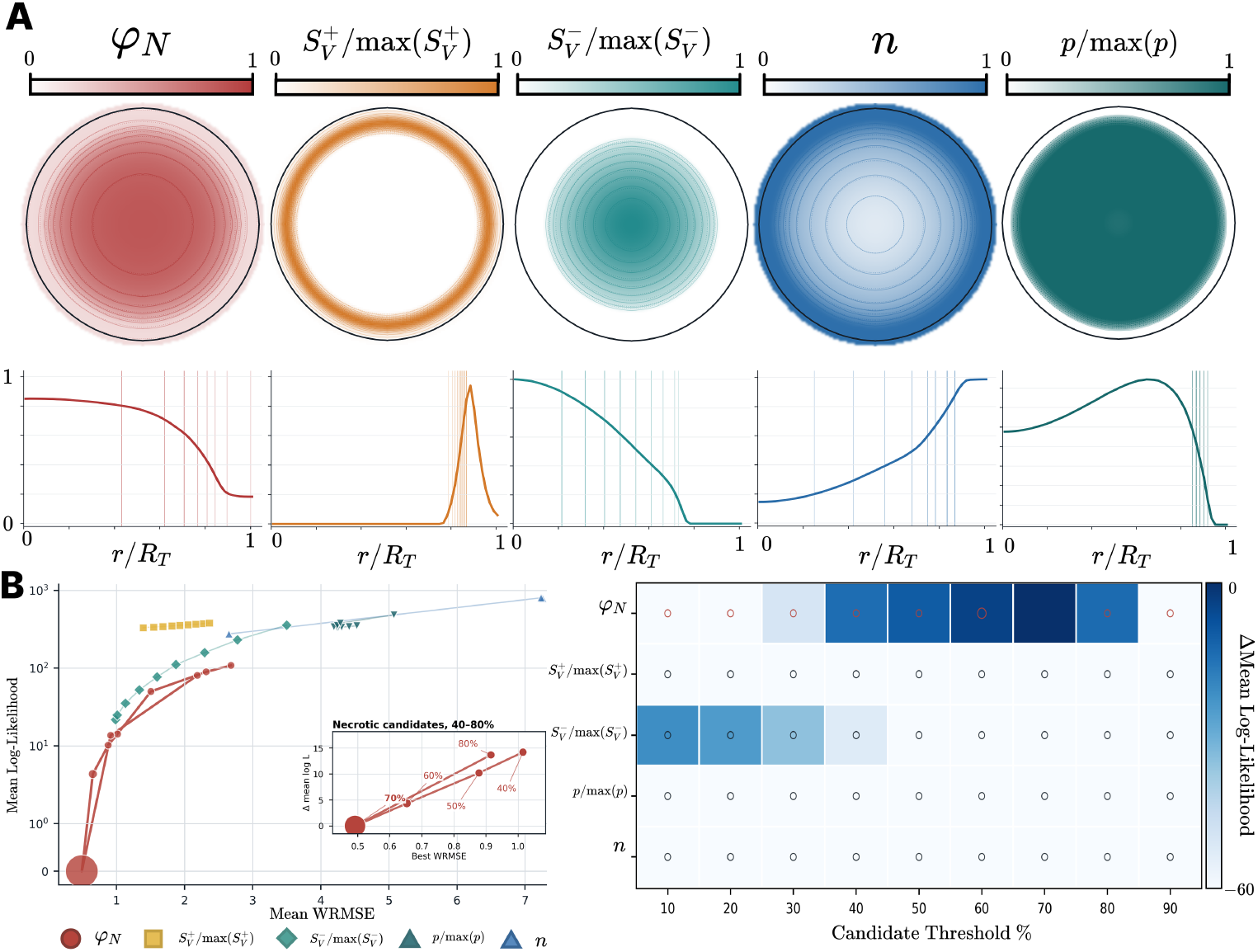
Exploratory comparison of continuum observables corresponding to the experimental inhibited radius. (A) Candidate intermediate-radius observables extracted from the final calibrated WM983 2.5k dataset (day 5). The upper panels show the necrotic-density (*φ*_*N*_), relative positive-source 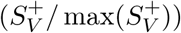, relative negative-source 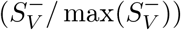, nutrient (*n*), and relative pressure (*p/* max(*p*)) fields. The lower panels show the corresponding averaged radial profiles; vertical lines identify the candidate thresholds evaluated for each observable family. (B) Representative exploratory all-candidate Bayesian SMC run for the WM983b 5k dataset under equal prior weight across candidate definitions. Left: candidate definitions evaluated during the run, displaying candidate families calibration performance. The strongest performing family (necrotic) is isolated in the focus panel, showing an optimum of the experimental objective (Eq. 12) within the candidate definitions. Right: relative exploratory support across candidate thresholds. The necrotic-structure family receives the strongest relative support in this example, with the 70% threshold receiving 98.06% weight of defining Browning’s inhibited radius.

**Figure 4:**
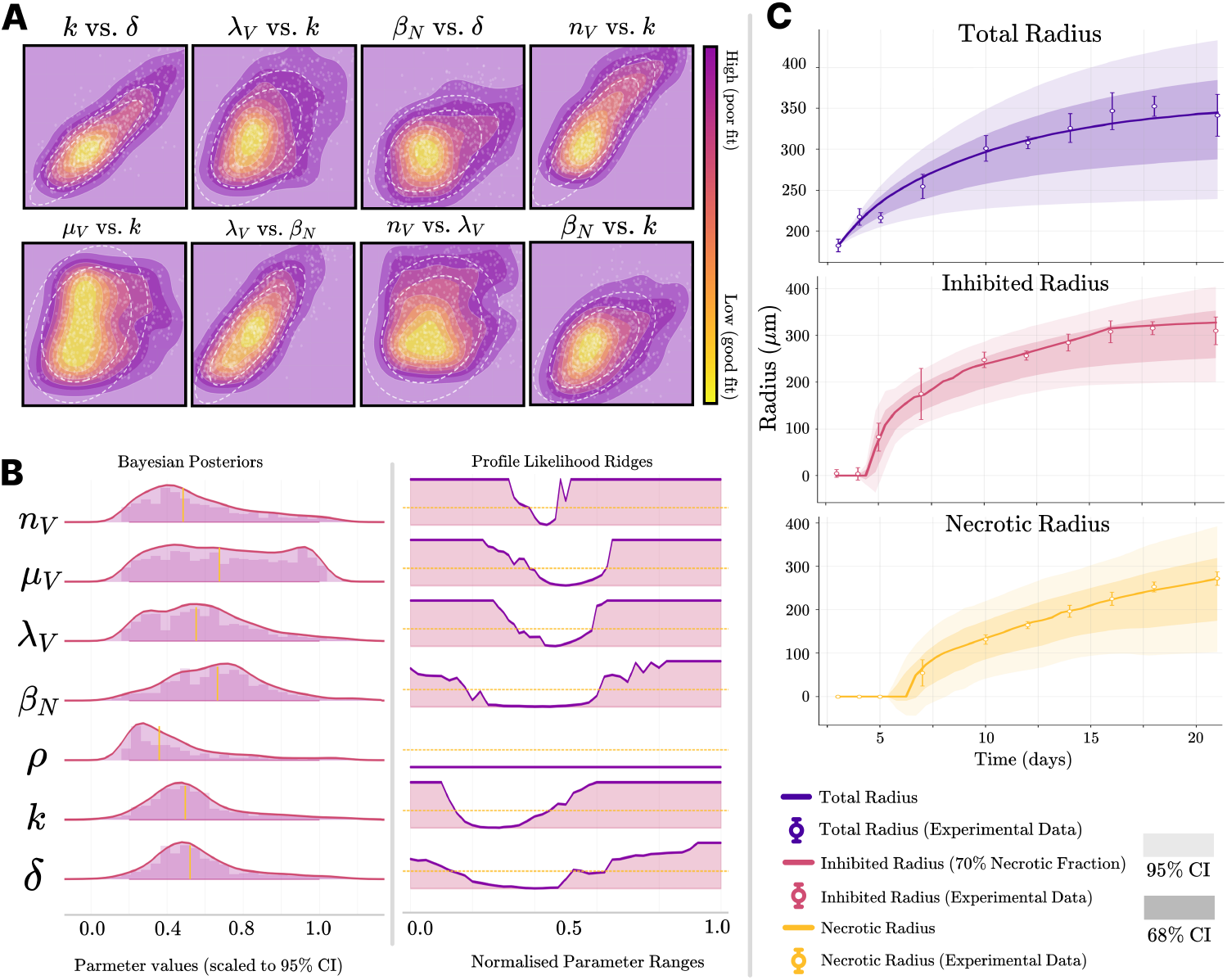
Posterior distributions from particle SMC for the WM983b 5k dataset. (A) SMC-identified basins in parameter-parameter space. The colour scale shows the fit quality, while the contour ellipses summarise posterior plausible regions. (B) Parameter posterior distributions (vertical line is posterior mean) and profile likelihood ridges (horizontal line identifies the 95% confidence interval baseline). (C) Simulated MAP versus the experimental data [37] showing total radius, inhibited radius, and necrotic core radius. The reported aggregate WRMSE is *C*(***θ***_MAP_) = 0.492.

**Figure 5:**
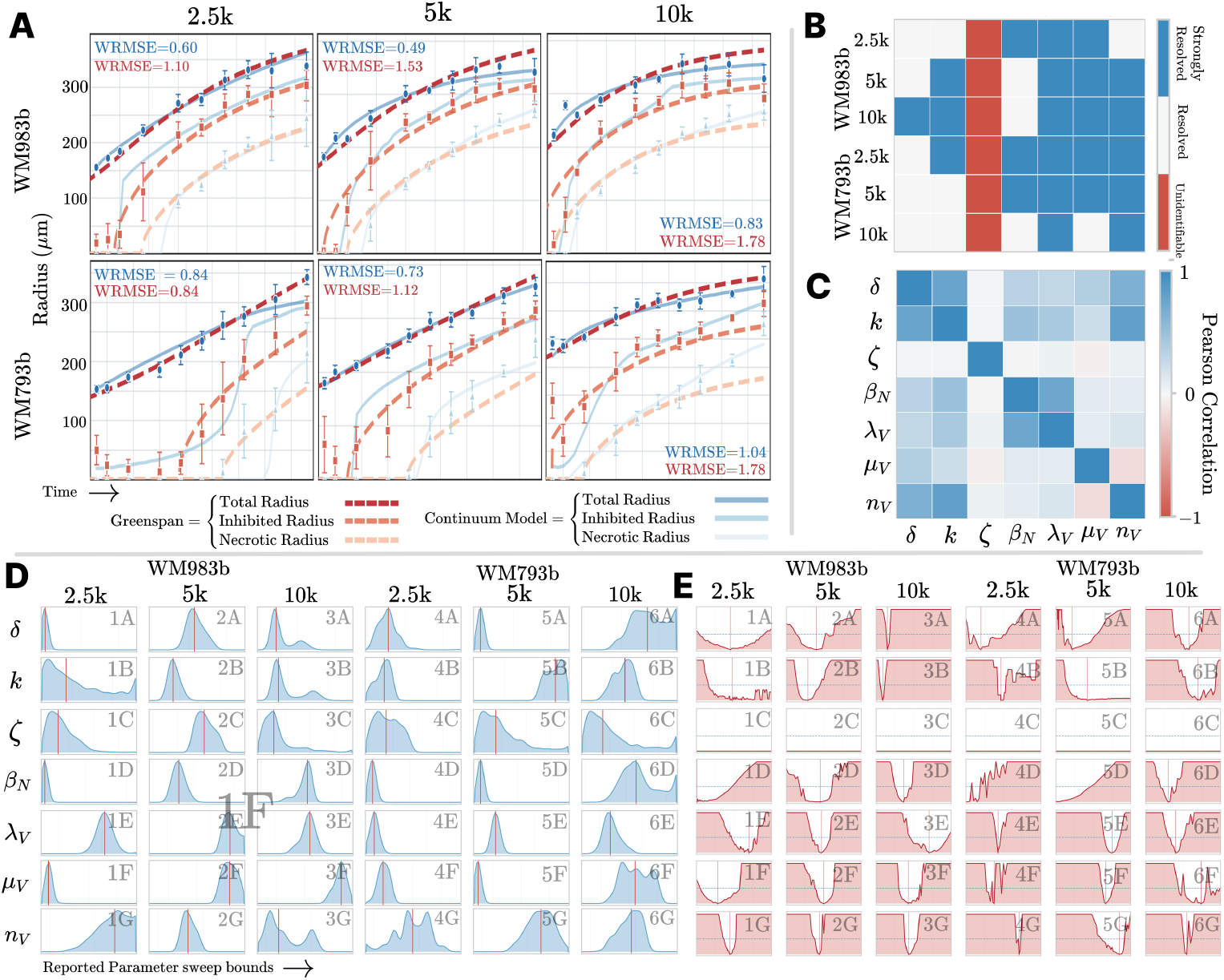
Full Dataset Calibration Results. (A) Experimental best fits for all calibrated datasets, compared to the Greenspan-type model. (B) Parameter identifiability heatmap across all calibrated datasets. (C) Combined parameter-parameter Pearson correlation metric across all calibrated datasets. (D) SMC posterior distributions for all parameters across all datasets, shown over the final refinement bounds. Vertical red line indicates the posterior mean. (E) Profile likelihood ridges for all parameters across all datasets, shown over the final refinement bounds. Horizontal shaded regions indicate the accepted refinement bounds.

### 5.1 Exploratory model-internal mapping of the inhibited radius

Across both the initial global exploration and the exploratory Bayesian SMC runs, multiple inhibited-radius candidates fit the experimental data well. The field-derived observables most aligned with the data were the necrotic structures, the relative proliferation and death magnitude structures, and the internal pressure structures, whose distinct radial definitions are shown in Figure 3(A). The global database revealed plausible starting regions and showed that different candidate families occupied distinct regions of parameter space. Exploratory SMC runs then compared the full candidate set under equal working weights. The necrotic structures received the strongest relative support across datasets, while the other structures did not generalise as consistently; the representative exploratory all-candidate comparison in Figure 3(B) shows this concentration explicitly. The most promising candidate-associated basin was then selected for each dataset and refined in a focused SMC run. This staged exploration and refinement was useful because each candidate definition could occupy its own basin in the objective landscape. The result should be interpreted narrowly: within the CTM, necrotic-structure proxies provide the strongest relative model-to-data agreement in this workflow. It does not establish that Browning’s experimental boundary is biologically necrotic. Across all datasets, the selected candidate varied slightly but was consistently found between the 50–70% necrotic-occupied radius. This identifies a testable model-internal mapping that could motivate future experiments using direct spatial markers, rather than a definitive interpretation of the experimental observable.

### 5.2 Identifiability and uncertainty

A key design principle of the present framework is the use of parameters that correspond directly to biologically interpretable cellular processes. Rates governing proliferation, death, nutrient consumption, and necrotic clearance are each associated with measurable biological behaviours and therefore provide a clear mechanistic interpretation of the calibrated model parameters. Within such continuum descriptions, however, it is important to emphasise that these quantities should be interpreted as effective parameters rather than direct microscopic constants [31, 30, 32]. Each calibrated parameter represents a coarse-grained summary of many underlying biological mechanisms, including intracellular regulatory networks, stochastic cell-fate decisions, and micro-environmental heterogeneity that cannot be resolved at the spatial and temporal scales of spheroid imaging data [21, 24]. Importantly, this effective-parameter interpretation should be viewed as a strength rather than a limitation of the approach. By operating at the continuum scale, the model avoids trying to resolve mechanisms that are neither experimentally observable nor practically identifiable from organoid-scale measurements, while still retaining a direct correspondence between model parameters and dominant biological processes [31, 32, 30]. This shift toward effective parameter interpretation aligns the model with replicate-style experiments. The use of replicates inherently acknowledges the stochastic nature of biological systems and reduces measurements on these systems to *effective* statistical averages, directly paralleling the interpretive framework of the continuum model.

Across all datasets, biological and nutrient-associated parameters were locally resolved within the accepted calibration basins, although some parameters were constrained through compensatory relationships, one-sided physical bounds, or saturation behaviour, as shown by the combined resolution, posterior, and profile-likelihood summaries in Figure 5(A,D,E). This result aligns with previous calibration attempts on this dataset across two different models [39, 37]. Both of these previous calibrations identify that model parameters are not constrained by the total radius alone, but once the inhibited and necrotic radius are included, model parameters become substantially better resolved. We achieve the same qualitative resolution pattern, and use the SMC posterior samples to further identify correlation structures within the model.

The nutrient-associated parameters *n*_*V*_ , *k, µ*_*V*_ , and *δ* form a compensatory manifold, where combinations of these parameters can trade off to produce similar-quality fits. This diffusion-limited identifiability structure is visible in the cross-dataset correlation and posterior summaries in Figure 5(B,D), and is closely aligned with many biological experiments that reveal the unidentifiability of parameters corresponding to nutrient diffusion and consumption through tissue. Parameter correlations also exist between *λ*_*V*_ and *β*_*N*_ , and between *n*_*V*_ and *µ*_*V*_. The correlation between an effective growth rate and an inhibitor of that growth rate is not unexpected, nor is the correlation between an effective death rate and a nutrient death threshold. Stronger growth rates tied to stronger inhibition could generate similar trajectories to weaker counterparts, and a similar compensatory structure exists between the effective death rate and the nutrient threshold. These correlation structures are consistent with well-documented “sloppiness” of biological mathematical models [59, 57], reflecting the limited information content of radius-based measurements alone.

The mechanical pressure constant *ζ* is the least biologically interpretable swept parameter in the current calibration. It affects the physical relaxation of the continuum fields, and is the only parameter that did not directly encode a cell fate process in the source terms. Its weak resolution is apparent in the cross-dataset resolution and profile-likelihood summaries in Figure 5(A,E). This suggests that the untreated radius data mostly constrain growth, nutrient response, and necrotic accumulation, while being less informative about mechanical pressure relaxation. This does not mean that *ζ* is fundamentally unidentifiable. In principle, it should become better resolved under experiments that perturb the mechanical environment, for example by growing spheroids in media, scaffold, or extracellular-matrix conditions that exert different levels of external stress, confinement, or resistance to expansion. This is consistent with experimental work showing that spheroid size, necrosis, viability, and structural integrity vary with oxygen, media composition, serum concentration, and matrix context, and with broader organoid engineering work that uses matrix design and mechano-physiological control to shape growth and reproducibility [18, 60, 4]. Under those conditions, the same growth and nutrient parameters would be observed under different mechanical loads, making the pressure-relaxation parameter more directly probed by the data. In later model versions this parameter may therefore either be fixed or narrowed for standard free-growth assays, or estimated explicitly under specific media-conditions, or scaffolds.

### 5.3 Comparison with the Greenspan-type model

The Greenspan-type model comparison remains an important baseline because it represents the canonical spheroid model. Its strength is its simplicity and direct alignment with classical spheroid structure. The core dynamic variables are themselves the canonical spheroid radii, and therefore the experimental data map directly onto the model compartments. This is also the limitation addressed by the continuum model. The Greenspan-type compartment topology, although highly aligned with untreated structure, cannot represent more complex spheroid structures that may appear under drug-treated conditions without structural reformulation. Under a simple assumption of strong viable cell death at high drug concentrations, the immediate consequence would be the formation of a necrotic annulus, introducing a fourth layer into the spheroid structure. Topologically, the Greenspan-type model cannot realise a state that induces a different ordered, layered structure such as this without changing its compartment structure. Because of the general field-based formulation of the continuum model, drug-treated conditions can be modelled through the addition of a quasi-steady drug concentration field. We provide a simple implementation of this in Section 4.4, with the resulting structural changes shown in Figure 6(B,C).

**Figure 6:**
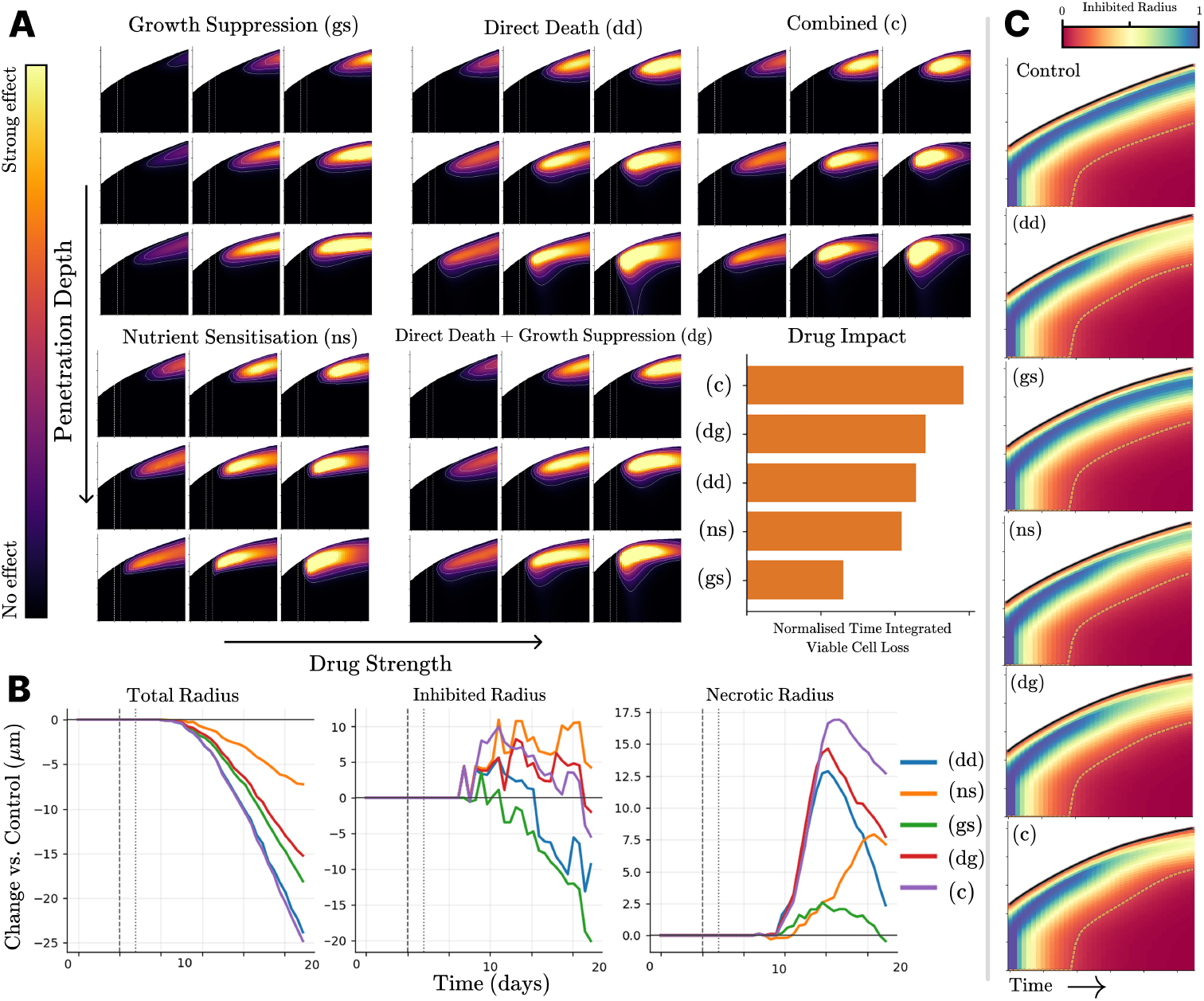
Drug-response field effects. (A) Radial profiles of drug-attributed viable cell loss across three strengths and penetration depths per drug-effect mechanism. Each panel represents the radial profile of a spheroid over a drugged simulation. For each 3 × 3 panel, left to right indicates increasing drug strength, and top to bottom indicates increasing characteristic penetration depth of the drug. (B) Variation of the canonical layered structure under drug-treated conditions. (C) Radial profiles of the viable cell population normalised to the final total radius. A diverging colourmap is used so the calibrated inhibited radius is implicitly visible as the divergence point of the colourmap.

The improvement in WRMSE for the continuum model therefore should be interpreted as more than a purely numerical fit improvement. The matched trajectories in Figure 5(C) show that the continuum model recovers the canonical three-region growth pattern with better agreement to the experimental data than a model whose structure is explicitly built around those three regions, while also being more flexible for modelling different external conditions. This supports the main modelling argument of the paper: the canonical compartment structure is still present and useful as a biological reference point, but does not need to be predefined into the model structure.

### 5.4 Limitations and future work

The present calibration is still limited by the observables available in the Browning dataset. Total radius, inhibited radius, and necrotic radius provide useful longitudinal constraints, but they are summary measurements of a richer spatial process. The use of a single threshold for the inhibited radius in the experimental data restricts the ability to use the full spatial profiling data that would have been available from raw images. This effectively provides only a single intermediate radius that can be calibrated to the data. This worked well for the Greenspan-type model, which only specifies one radius for that region, but it disregards much of the information about internal gradients and structures throughout the spheroid. The model formulation, and the interior profiling infrastructure illustrated in Figure 3(A), allows entire continuum radial profiles to be calibrated, making use of information left unavailable by the reduced Browning *et al*. [37] measurements. Thresholding the continuum radial profiles into incrementally smaller partitions leaves room to expand the information density used for calibration. Further experiments for stronger patient calibration could use this by calibrating many thresholded intermediate radii across different observable definitions. Furthermore, by moving away from the classical *inhibited radius*, we can provide stronger simulation-to-experiment functional mappings between observables. These measurements could couple to produce calibration regimes that calibrate total radius, necrotic core radius, and full intermediate radial profiles, substantially increasing the capacity for data calibration compared to the calibration applied in this paper.

More generally, the model should not be limited to the radial growth experiment considered here. Because the governing equations are defined over continuous fields, different experimental configurations can be represented by changing the initial and boundary conditions rather than changing the model structure. Spheroid fusion experiments are a natural example. During these experiments, two or more spheroids are brought into contact and the rate at which they merge is controlled by the balance of cell-cell adhesion, interfacial energy, motility, and mechanical relaxation [61, 62]. This would provide a more direct way to probe the energy scale of the adhesion parameters than radius-only growth data, and could reveal whether different patients, cancer types, or culture conditions generate different effective cohesive strengths. This is also biologically meaningful for spheroid formation itself. Some cell types form compact spheroids readily, while others produce loose aggregates or fail to form stable spheroids, and a continuum model with an explicit adhesion energy scale provides a way to ask whether those differences are consistent with weaker tissue cohesion, or stronger dispersal [47, 63, 22].

## 6 Conclusion

This work demonstrates that a general continuum phase-field model can recover the classical three-region structure of untreated spheroid growth without prescribing that structure in the model geometry. The calibrated model fits the Browning spheroid datasets [37] stronger on average by 40% to the comparison Greenspan-type model [39], while also comparing candidate continuum observables as possible model-internal proxies for the experimentally measured inhibited radius. Across datasets, and under the exploratory candidate weighting used here, necrotic-structure proxies received the strongest relative support.

The calibration shows local resolution of most model parameters within the accepted calibration basins, with the pressure parameter *ζ* remaining weakly resolved. The biological and nutrient-associated parameters are reportable within the accepted basins, but their interpretation is not one-dimensional. Nutrient sensitivity, nutrient threshold, diffusion hindrance, cell death, and necrotic feedback form compensatory structures that describe effective tissue-scale behaviours, and some parameters are supported through boundary or saturation regimes rather than symmetric interior optima. The untreated calibration provides the baseline model validity required for the broader aim of this framework. The model’s flexibility under drug-treated conditions is demonstrated as a proof of concept for further exploration.

## Author Contributions

**Riley J. McNamara**: Conceptualization, Methodology, Software, Validation, Formal Analysis, Investigation, Data Curation, Writing – original draft, Visualization. **Mark C. Allenby**: Supervision, Writing – review and editing. **Gloria M. Monsalve-Bravo**: Supervision, Writing – review and editing. **Sandra R. Stein**: Writing – review and editing. **Glenn D. Francis**: Writing – review and editing.

## Data Availability

The experimental data used for calibration are publicly available from Browning et al. [37]. The simulation code and derived calibration outputs are not currently deposited in a public repository because their release is subject to approval from Genomics for Life Pty Ltd. They are available from the corresponding author upon reasonable request, subject to that approval.

## Supplementary Information

Supplementary Information contains the model derivation, nondimensionalisation, calibration database description, reportable parameter tables, experimental design, drug-response source-term details, and Bayesian parameter-pair distributions for all six calibrated datasets. It is submitted as a single separate file with the manuscript.

## Declaration of Generative AI and AI-Assisted Technologies in the Manuscript Preparation Process

During the preparation of this work, the authors used GPT-5.6-Sol to proofread, review, build L^A^TEX documents, and resolve BibTeX references. After using these tools, the authors reviewed and edited the content as needed and take full responsibility for the content of the published article.

## Acknowledgements

The authors acknowledge the support of the University of Queensland and Genomics for Life Pty Ltd.

## Supplementary Information

This file contains the supporting derivations, calibration details, parameter reports, drug-response source-term formulation, and Bayesian parameter-pair distributions for the main manuscript.

## S1 Phenotypic Conversion

Mass transfer between populations—representing phenotypic drift, infection-like processes, or differentiation—is handled by modifying the proliferation term. Let *p*_*i*_ denote the fraction of new growth in population *i* that remains in *i*. The remaining fraction (1 − *p*_*i*_) is transferred to another population *j*. This yields

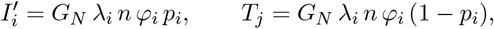

so that 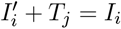 and mass is conserved.

### Generalised transfer

For models with more than two interacting phenotypes, we introduce a full set of transfer fractions *p*_*i*→*k*_ satisfying

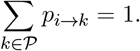

Incoming mass for population *i* becomes 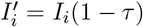 with *τ* = 1 − *p*_*i*→*i*_, and each recipient population *k* ≠ *i* obtains a transfer term *T*_*k*_ = *I*_*i*_ *p*_*i*→*k*_.

### S1.1 Vector Formulation

To streamline implementation and generalisation, while unlocking hardware-accelerated linear-algebra kernels, we collect all *M* cell-phase fields into a (1 *× M*) vector:

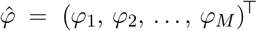

Note that we fix the necrotic population to index *M* such that *φ*_*M*_ := *φ*_*N*_. We then introduce likewise-stacked mass fluxes 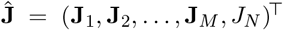. , and solid velocities 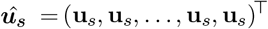.

The phase-field evolution equation becomes the compact system:

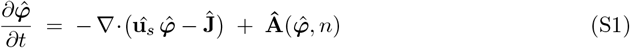

where ***A*** is the generalised source vector whose entries encode phenotype-specific proliferation, death, and conversion rates (including the smooth nutrient switches and the global necrotic feedback *G*_*N*_):

To describe **A**, we first define the transfer matrix **P** as:

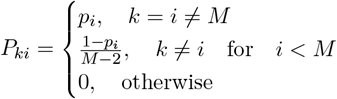

Such that the sum of each column is: 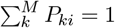. We then define 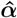, whose elements are *α*_*i*_ = *λ*_*i*_*φ*_*i*_, and ***D*** as the “death matrix”, whose elements are:

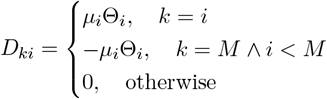

We can now define **A** as:

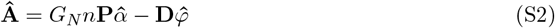

For a Healthy (H), Diseased (D), and Necrotic (N) triad the structure is

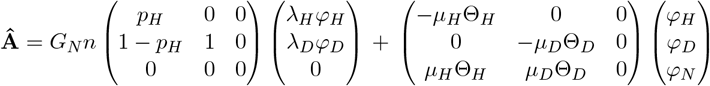

This formulation recapitulates any model with *M* different cell populations into a single equation, allowing all terms to be computed through unified vectorised operations. In the present study, necrotic clearance is omitted by setting *µ*_*N*_ = 0. This structure enables seamless extensibility; new populations can be incorporated by appending entries to these vectors, allowing efficient implementation and execution via parallelised linear algebra kernels.

## S2 Objective Functional Form

We define *Ô*_sim_(*t*_*i*_; ***θ***) = (*R*_T,sim_, *R*_I,sim_, *R*_N,sim_) and *Ô*_exp_(*t*_*i*_) = (*R*_T,exp_, *R*_I,exp_, *R*_N,exp_) with corresponding experimental standard errors 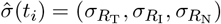. The WRMSE objective is:

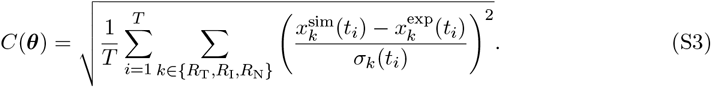

Here the denominator is *T* rather than *T* |*O*| because the reported score uses the aggregate standardised residual across the three radii at each time point. Thus *C* is not the RMS per individual observation; the corresponding per-observation RMS is 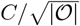.

## S3 Model Nondimensionalisation

Experimental spheroid datasets span a range of initial seeding densities and cell lines, each producing a different initial spheroid radius *R*_0_. A simulation library generated directly in physical coordinates would therefore require separate simulations for each initial size. We instead non-dimensionalise the model using the dataset-specific initial radius. This gives a general radius-scaled model first. We then introduce a single sweep parameter that records how the fixed nutrient penetration depth compares with the radius represented by a given simulation.

### S3.1 Characteristic scales from the nutrient field

The relevant transport length is set by the balance between nutrient transport and consumption. Ignoring density-dependent hindrance for the moment, a one-dimensional steady-state balance has the form:

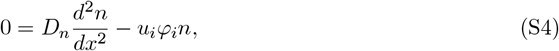

where *D*_*n*_ is the baseline external-medium diffusivity and *u*_*i*_ is the consumption rate for population *i*. Dimensional balancing gives the familiar penetration-depth scale:

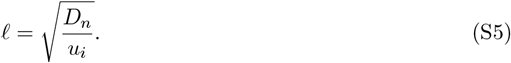

In the full model, *ℓ* should be interpreted as an effective penetration depth rather than as a direct estimate of *u*_*V*_. The hindrance function modifies transport inside dense tissue, so this characteristic length reflects the combined effect of baseline diffusivity, cellular uptake, and density-dependent transport hindrance. We treat *ℓ* as a fixed characteristic length for a given cancer type or experimental condition, chosen from literature or prior calibration, while *D*_*n*_ remains the baseline external-medium diffusivity. The coordinate scale used in the simulation is not *ℓ* itself, but the dataset-specific initial radius *R*_0_.

### S3.2 Radius-based scaling

With *R*_0_ and *T*_0_ defined, we introduce non-dimensional coordinates, differential operators, and fields:

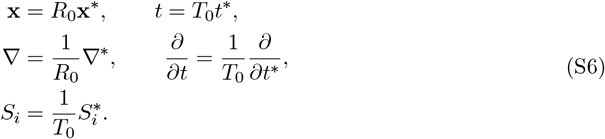

The cell volume fractions *φ*_*i*_ and nutrient concentration *n* are already dimensionless, so no additional field scaling is required for them. The pressure scaling is introduced explicitly below when deriving the pressure equation. The initial spheroid radius is fixed at 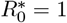 in simulation units, and physical distances are recovered by multiplying dimensionless distances by *R*_0_.

### S3.3 Nutrient equation

Let

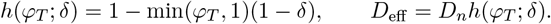

Using **x** = *R*_0_**x**^∗^ gives

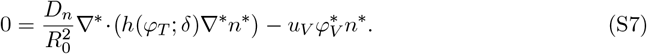

Dividing by 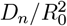 and using *ℓ*^2^ = *D*_*n*_*/u*_*V*_ yields

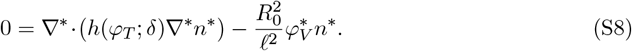

We note that the nutrient equation no longer relies on only the non-dimensional length scale, but the ratio of that length scale to the physical penetration depth.

### S3.4 Free energy

The Cahn–Hilliard free energy functional:

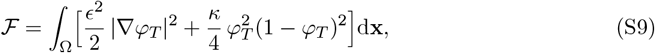

has units of energy. To make the energetic part of the model fully non-dimensional, we first factor out the bulk free-energy density scale *κ* and define the physical interface-width scale

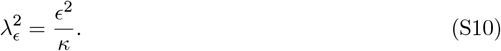

Then

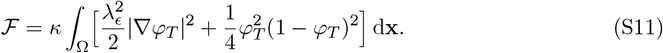

Substituting **x** = *R*_0_**x**^∗^ gives

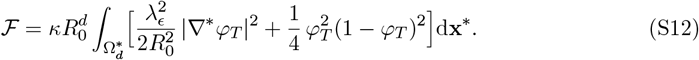

Choosing the energy scale 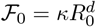 gives the dimensionless free energy

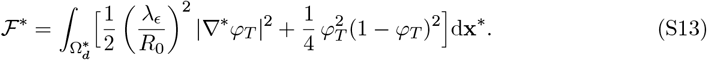

Thus the physical interface width enters the non-dimensional model through the radius-scaled group 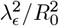.

### S3.5 Chemical potential

The chemical potential is the variational derivative of the free energy:

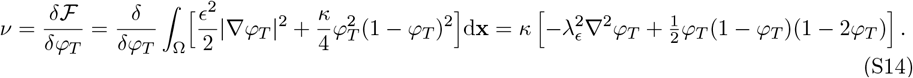

Scaling the chemical potential by *ν* = *κν*^∗^ and substituting ∇ = ∇^∗^*/R*_0_ gives

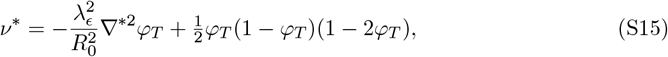

where 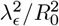 is the corresponding dimensionless interface-width group. The diffusive flux **J**_*i*_ = *m*_*i*_∇*ν* can then be scaled consistently with the chosen time and chemical-potential scales:

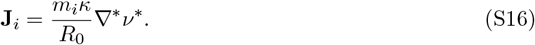

### S3.6 Pressure Poisson equation

The pressure satisfies:

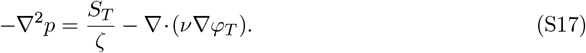

Substituting ∇ = ∇^∗^*/R*_0_ and 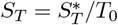:

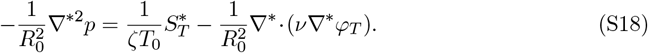

Multiplying through by 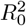 and defining *p*^∗^ = *p* yields:

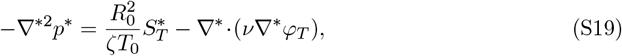

where 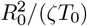 is the corresponding dimensionless source-pressure group. In the collected system below, pressure is scaled so that this group is absorbed into the definition of the dimensionless pressure and source terms.

### S3.7 Solid velocity

The Darcy-like solid velocity:

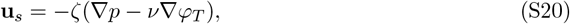

becomes in non-dimensional form:

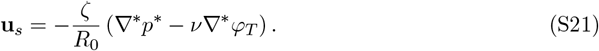

### S3.8 Source terms

The local source terms are already defined in non-dimensional form in the main text. The proliferation rate *λ*_*V*_ and viable-cell death rate *µ*_*V*_ carry units of 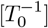, and the remaining local parameters (*n*_*V*_ , *δ, ζ, k*) are dimensionless or scaled consistently. Necrotic clearance is set to *µ*_*N*_ = 0 throughout this study.

The global necrotic feedback requires one additional scaling because it contains an integral over space. In *d* spatial dimensions,

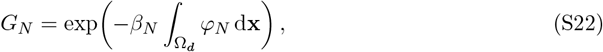

so *β*_*N*_ has units of length^−*d*^. Under the radius-based scaling **x** = *R*_0_**x**^∗^,

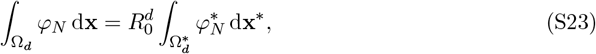

and therefore

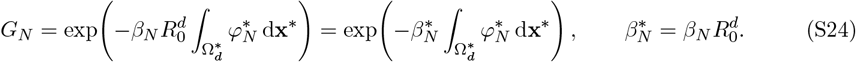

Thus the parameter appearing in the non-dimensional simulation is the dimensionless feedback strength 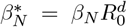. If reporting a physical feedback strength, 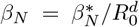, for two-dimensional simulations this has units of m^−2^.

### S3.9 Introducing the sweep parameter

The radius-based system above contains *R*_0_ through three length-dependent groups:

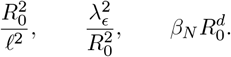

To generate one dimensionless simulation library that can represent multiple initial spheroid sizes, we introduce

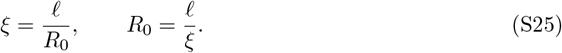

For fixed *ℓ*, sweeping *ξ* is equivalent to sweeping the initial physical radius represented by the dimensionless simulation. The coordinate *ξ* is therefore a library index used to cover the different initial-radius regimes represented by the six datasets; it is included in global sweeps and scoring, but is not interpreted as an additional biological mechanism or as an independently reportable cell-fate parameter. The substitutions are

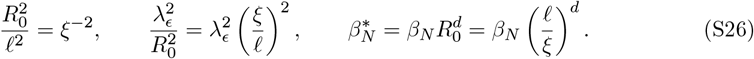

For a new dataset, *R*_0_ is measured from the initial spheroid image and the matching library slice is identified through *ξ* = *ℓ/R*_0_. The surrounding sweep values are retained in the universal database so that the dataset-specific comparison can select the appropriate scale regime. After the simulation has been selected at that *ξ*, any simulated distance is returned to physical units using

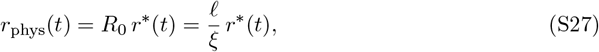

where *r*^∗^(*t*) may denote the total radius, necrotic radius, inhibited radius, or any other distance extracted from the non-dimensional fields.

### S3.10 Fully non-dimensionalised system

After substituting Eq. (S26) and omitting asterisks for readability, the sweep-indexed system solved in simulation is:

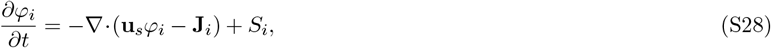

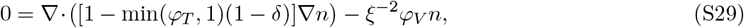

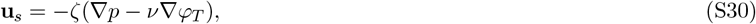

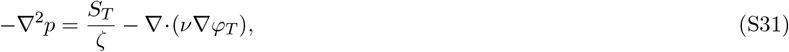

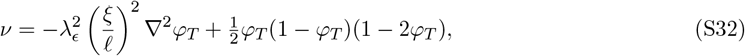

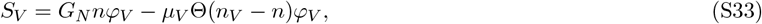

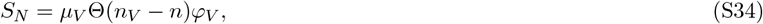

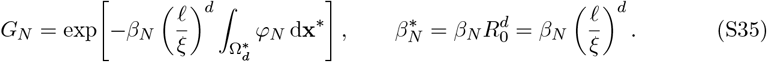

This system is solved on the unit-order domain Ω^∗^ with *n* = 1 on *∂*Ω^∗^ and homogeneous Neumann conditions on *φ*_*i*_ and *p*.

### S3.11 Physical consistency across different initial radii

Consider two datasets from the same cancer type, cultured in the same external medium, with initial radii *R*_1_ and *R*_2_. They share the same physical *D*_*n*_, the same characteristic penetration depth *ℓ*, the same physical interface width *λ*_*ϵ*_, and the same physical feedback strength *β*_*N*_ , but they have different sweep values

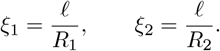

For dataset *j*, the nutrient equation in the simulation is

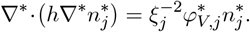

Mapping this term back to dimensional coordinates gives

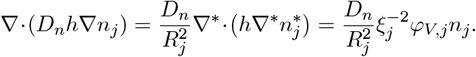

Since *ξ*_*j*_ = *ℓ/R*_*j*_,

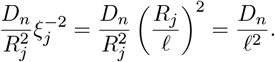

Thus changing *R*_*j*_ changes the dimensionless coefficient in the simulation, but it does not create a new physical diffusivity. The external-medium diffusivity remains the same dimensional quantity *D*_*n*_ with units of length squared per time.

The interface-width scaling is consistent in the same way. For dataset *j*, the dimensionless coefficient in the chemical potential is

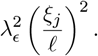

Mapping the gradient term back to physical coordinates gives

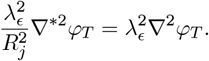

Thus each simulation uses a different dimensionless interface width *λ*_*ϵ*_*/R*_*j*_, but this corresponds to the same physical interface width 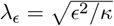.

The same logic applies to the necrotic feedback parameter. For dataset *j*,

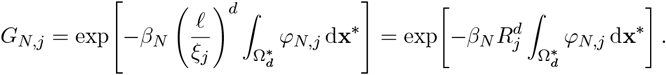

Using 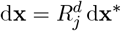, this is exactly

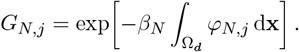

Therefore the dimensionless coefficient changes with the represented radius, 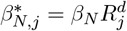, but the physical parameter *β*_*N*_ retains the same units and meaning. In two-dimensional simulations *d* = 2, so *β*_*N*_ has units of m^−2^; in three-dimensional simulations *d* = 3, it has units of m^−3^.

## S4 Simulation Library and Comparison Database

All simulation outputs and experimental comparisons are stored in a DuckDB-backed Parquet database with two layers. The *Universal Library* records nondimensional simulation runs indexed by the sweep coordinate *ξ* (Supplementary Information Section S3), making outputs reusable across any experimental dataset. The *Comparison Database* stores scored comparisons between library runs and individual datasets, including per-run WRMSE values as defined in (S3).

### Universal Library

Each sweep (`sweeps` table) spawns many runs (`runs`), keyed by *ξ* and *χ* = *ξ*^−2^. Per-run outputs are stored in three tidy-format tables: `observables` (time-series scalar radii such as Total_Radius, Necrotic_Radius, and multiple interior candidate columns), `observable_profiles` (radial profiles of any field), and `parameters` (sampled parameter values). A `run_behavior_descriptors` table records cheap diagnostics—growth classification, necrotic core presence, and anomaly flags—used to filter runs before scoring (e.g., excluding boundary-contact or numerically unstable runs).

### Comparison Database

Experimental datasets (`datasets` table) provide initial radius *R*_0_, penetration depth *ℓ*, reference timescale *T*_ref_, and per-timepoint observations with uncertainties (`dataset_observations`). Each dataset–sweep pair is tracked by a `comparison_jobs` entry with a hash-based identifier and version fingerprints for change detection. Scoring produces two result tables: `comparison_run_scores` records the best candidate column and combined WRMSE per run, while `comparison_candidate_scores` logs all candidate-level diagnostics with per-candidate RMSE and rank.

### Scoring pipeline

Experimental observables are mapped from dimensional units (µm, days) to dimensionless simulation coordinates using the dataset-specific scales *R*_0_ and *T*_ref_. Each run produces multiple candidate columns for interior observables (e.g., isodensity radii at different thresholds); all candidates are evaluated against each experimental target. The best candidate per run is selected by minimising the combined WRMSE. Behavior filters from `run_behavior_descriptors` exclude anomalous runs before scoring.

The global exploratory sweep used the inclusive parameter bounds listed in Table S1. These bounds define the broad search domain before the accepted basin and profile-likelihood refinements described below.

**Table S1:**
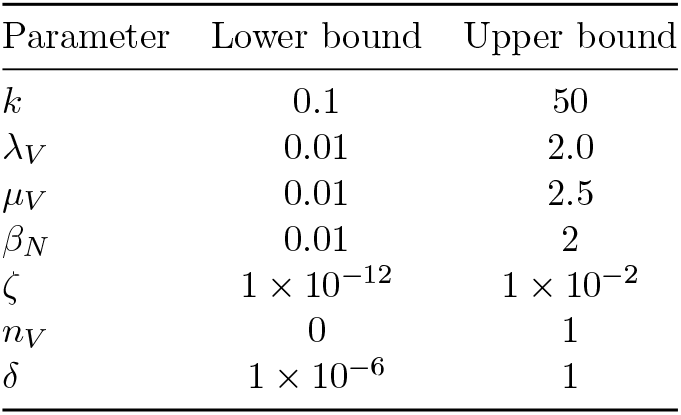
Global parameter bounds used for the exploratory sweep of the continuum model.

| Parameter | Lower bound | Upper bound |
| --- | --- | --- |
| $k$ | 0.1 | 50 |
| $\lambda_V$ | 0.01 | 2.0 |
| $\mu_V$ | 0.01 | 2.5 |
| $\beta_N$ | 0.01 | 2 |
| $\zeta$ | $1 \times 10^{-12}$ | $1 \times 10^{-2}$ |
| $n_V$ | 0 | 1 |
| $\delta$ | $1 \times 10^{-6}$ | 1 |

## S5 Reportable Browning Fit Parameters

**Table S2:**
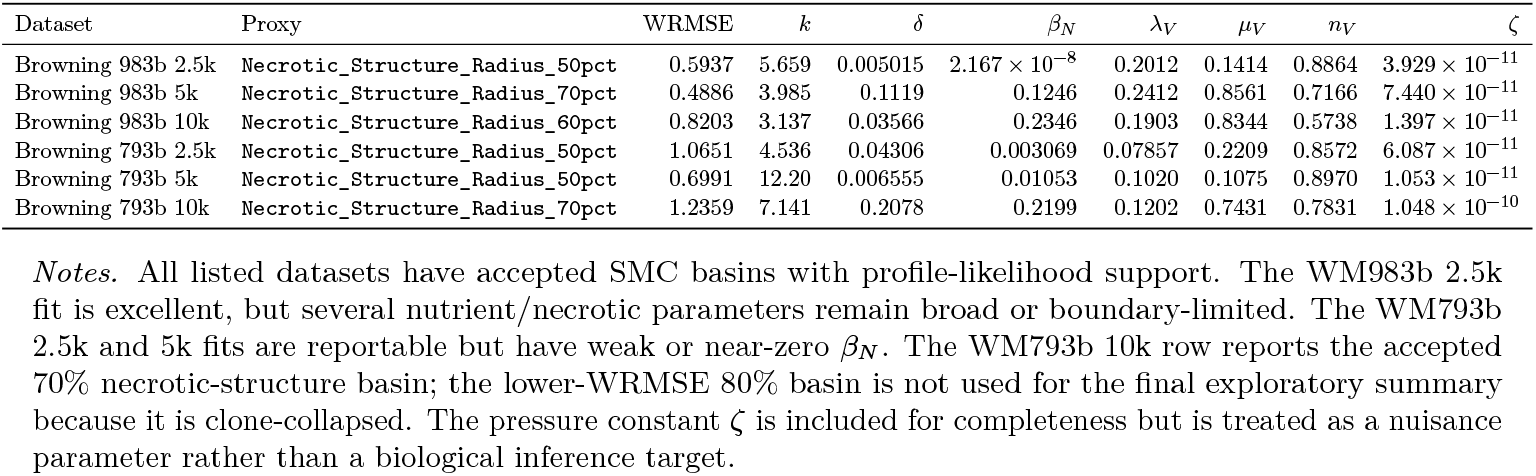
Currently reportable Browning best-fit parameter values from the accepted SMC/profile-likelihood calibration basins. WRMSE values correspond to the reportable profile-supported fits where available.

| Dataset | Proxy | WRMSE | $k$ | $\delta$ | $\beta_N$ | $\lambda_V$ | $\mu_V$ | $n_V$ | $\zeta$ |
| --- | --- | --- | --- | --- | --- | --- | --- | --- | --- |
| Browning 983b 2.5k | Necrotic_Structure_Radius_50pct | 0.5937 | 5.659 | 0.005015 | $2.167 \times 10^{-8}$ | 0.2012 | 0.1414 | 0.8864 | $3.929 \times 10^{-11}$ |
| Browning 983b 5k | Necrotic_Structure_Radius_70pct | 0.4886 | 3.985 | 0.1119 | 0.1246 | 0.2412 | 0.8561 | 0.7166 | $7.440 \times 10^{-11}$ |
| Browning 983b 10k | Necrotic_Structure_Radius_60pct | 0.8203 | 3.137 | 0.03566 | 0.2346 | 0.1903 | 0.8344 | 0.5738 | $1.397 \times 10^{-11}$ |
| Browning 793b 2.5k | Necrotic_Structure_Radius_50pct | 1.0651 | 4.536 | 0.04306 | 0.003069 | 0.07857 | 0.2209 | 0.8572 | $6.087 \times 10^{-11}$ |
| Browning 793b 5k | Necrotic_Structure_Radius_50pct | 0.6991 | 12.20 | 0.006555 | 0.01053 | 0.1020 | 0.1075 | 0.8970 | $1.053 \times 10^{-11}$ |
| Browning 793b 10k | Necrotic_Structure_Radius_70pct | 1.2359 | 7.141 | 0.2078 | 0.2199 | 0.1202 | 0.7431 | 0.7831 | $1.048 \times 10^{-10}$ |

*Notes.* All listed datasets have accepted SMC basins with profile-likelihood support. The WM983b 2.5k fit is excellent, but several nutrient/necrotic parameters remain broad or boundary-limited. The WM793b 2.5k and 5k fits are reportable but have weak or near-zero $\beta_N$ . The WM793b 10k row reports the accepted 70% necrotic-structure basin; the lower-WRMSE 80% basin is not used for the final exploratory summary because it is clone-collapsed. The pressure constant $\zeta$ is included for completeness but is treated as a nuisance parameter rather than a biological inference target.

## S6 Experimental Design

We take standard values of *κ* and *ϵ* from the literature [1, 2]. Cell motility is fixed to a mid-range literature value, on the order of 10^−10^[cm^2^*/*s] [3, 4]. Necrotic clearance is omitted from this model by setting *µ*_*N*_ = 0, because necrotic dissolution is negligible over the spheroid experimental time scales and has no measurable effect on the observable radii.

where *λ*_*V*_ is the viable-cell proliferation rate [t^−1^], *µ*_*V*_ the nutrient-dependent death rate [t^−1^], *n*_*V*_ the dimensionless nutrient satiety threshold, *δ* the nutrient diffusion hindrance [unitless], *ζ* the mechanical pressure constant [unitless], *β*_*N*_ the global necrotic-feedback strength [m^−*d*^] (with *d* = 2 for two-dimensional simulations), *k* the steepness of the nutrient switch [unitless], and *ξ* the ratio of penetration depth to initial radius [unitless].

## S7 Parameter Report

**Table S3:**
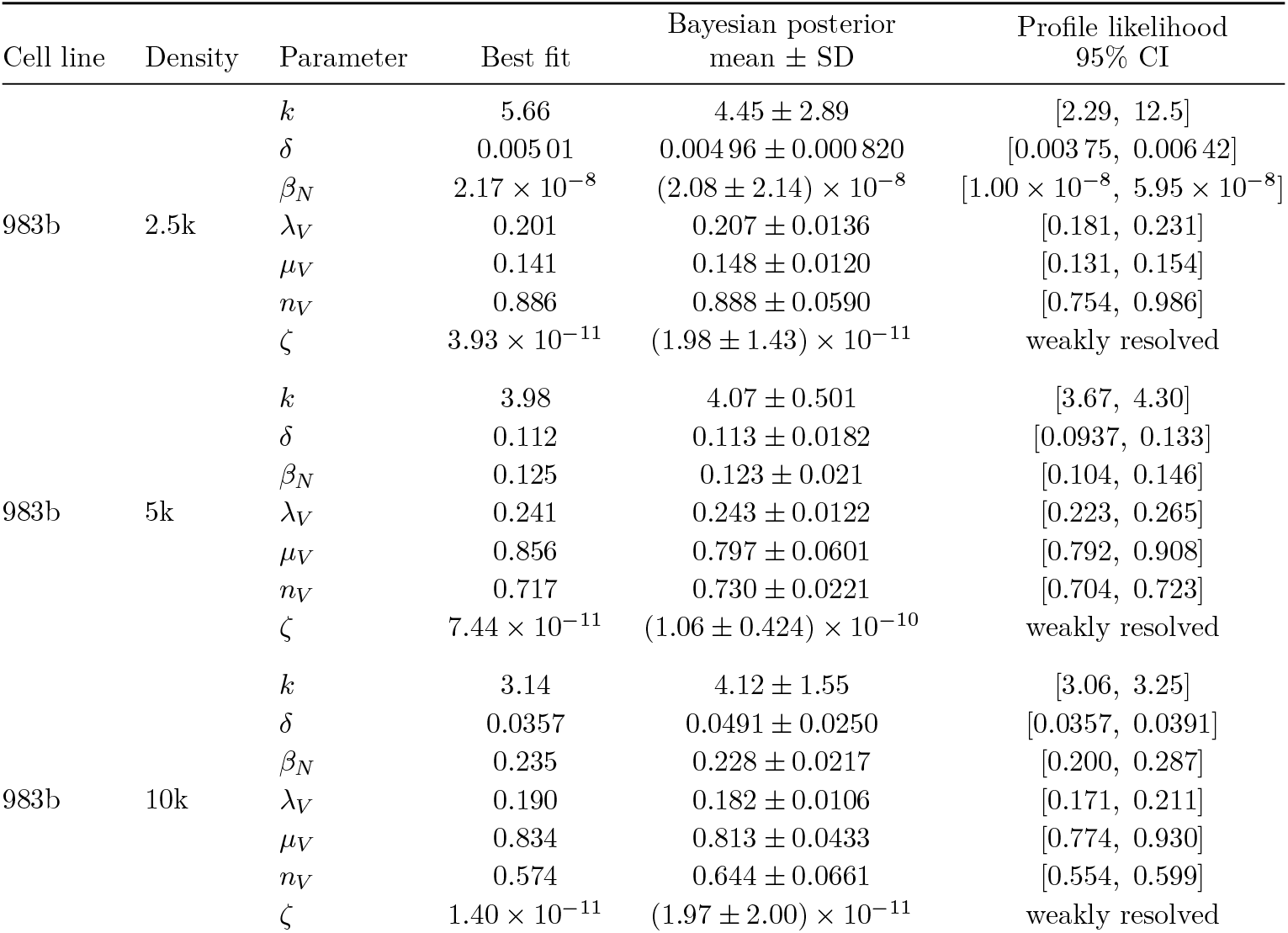
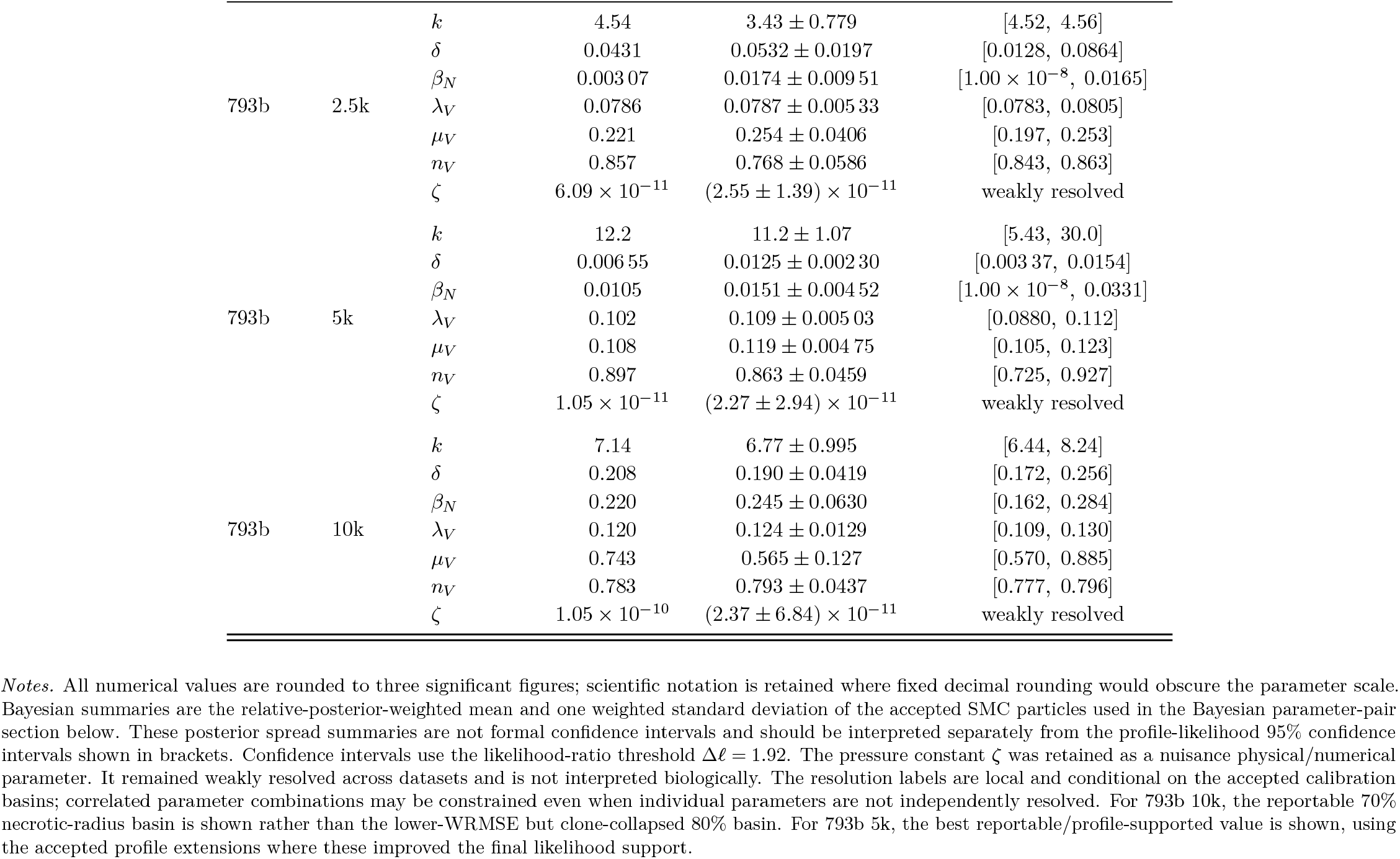
Best-fit calibrated parameters, weighted Bayesian posterior summaries, and profile-likelihood confidence intervals for the Browning spheroid datasets.

| Cell line | Density | Parameter | Best fit | Bayesian posterior<br>mean ± SD | Profile likelihood<br>95% CI |
| --- | --- | --- | --- | --- | --- |
| 983b | 2.5k | $k$ | 5.66 | $4.45 \pm 2.89$ | [2.29, 12.5] |
| | | $\delta$ | 0.005 01 | $0.004\,96 \pm 0.000\,820$ | [0.003 75, 0.006 42] |
| | | $\beta_N$ | $2.17 \times 10^{-8}$ | $(2.08 \pm 2.14) \times 10^{-8}$ | $[1.00 \times 10^{-8}, 5.95 \times 10^{-8}]$ |
| | | $\lambda_V$ | 0.201 | $0.207 \pm 0.0136$ | [0.181, 0.231] |
| | | $\mu_V$ | 0.141 | $0.148 \pm 0.0120$ | [0.131, 0.154] |
| | | $n_V$ | 0.886 | $0.888 \pm 0.0590$ | [0.754, 0.986] |
| | | $\zeta$ | $3.93 \times 10^{-11}$ | $(1.98 \pm 1.43) \times 10^{-11}$ | weakly resolved |
| 983b | 5k | $k$ | 3.98 | $4.07 \pm 0.501$ | [3.67, 4.30] |
| | | $\delta$ | 0.112 | $0.113 \pm 0.0182$ | [0.0937, 0.133] |
| | | $\beta_N$ | 0.125 | $0.123 \pm 0.021$ | [0.104, 0.146] |
| | | $\lambda_V$ | 0.241 | $0.243 \pm 0.0122$ | [0.223, 0.265] |
| | | $\mu_V$ | 0.856 | $0.797 \pm 0.0601$ | [0.792, 0.908] |
| | | $n_V$ | 0.717 | $0.730 \pm 0.0221$ | [0.704, 0.723] |
| | | $\zeta$ | $7.44 \times 10^{-11}$ | $(1.06 \pm 0.424) \times 10^{-10}$ | weakly resolved |
| 983b | 10k | $k$ | 3.14 | $4.12 \pm 1.55$ | [3.06, 3.25] |
| | | $\delta$ | 0.0357 | $0.0491 \pm 0.0250$ | [0.0357, 0.0391] |
| | | $\beta_N$ | 0.235 | $0.228 \pm 0.0217$ | [0.200, 0.287] |
| | | $\lambda_V$ | 0.190 | $0.182 \pm 0.0106$ | [0.171, 0.211] |
| | | $\mu_V$ | 0.834 | $0.813 \pm 0.0433$ | [0.774, 0.930] |
| | | $n_V$ | 0.574 | $0.644 \pm 0.0661$ | [0.554, 0.599] |
| | | $\zeta$ | $1.40 \times 10^{-11}$ | $(1.97 \pm 2.00) \times 10^{-11}$ | weakly resolved |
| 793b | 2.5k | $k$ | 4.54 | $3.43 \pm 0.779$ | [4.52, 4.56] |
| | | $\delta$ | 0.0431 | $0.0532 \pm 0.0197$ | [0.0128, 0.0864] |
| | | $\beta_N$ | 0.003 07 | $0.0174 \pm 0.009 51$ | $[1.00 \times 10^{-8}, 0.0165]$ |
| | | $\lambda_V$ | 0.0786 | $0.0787 \pm 0.005 33$ | [0.0783, 0.0805] |
| | | $\mu_V$ | 0.221 | $0.254 \pm 0.0406$ | [0.197, 0.253] |
| | | $n_V$ | 0.857 | $0.768 \pm 0.0586$ | [0.843, 0.863] |
| | | $\zeta$ | $6.09 \times 10^{-11}$ | $(2.55 \pm 1.39) \times 10^{-11}$ | weakly resolved |
| 793b | 5k | $k$ | 12.2 | $11.2 \pm 1.07$ | [5.43, 30.0] |
| | | $\delta$ | 0.006 55 | $0.0125 \pm 0.002 30$ | [0.003 37, 0.0154] |
| | | $\beta_N$ | 0.0105 | $0.0151 \pm 0.004 52$ | $[1.00 \times 10^{-8}, 0.0331]$ |
| | | $\lambda_V$ | 0.102 | $0.109 \pm 0.005 03$ | [0.0880, 0.112] |
| | | $\mu_V$ | 0.108 | $0.119 \pm 0.004 75$ | [0.105, 0.123] |
| | | $n_V$ | 0.897 | $0.863 \pm 0.0459$ | [0.725, 0.927] |
| | | $\zeta$ | $1.05 \times 10^{-11}$ | $(2.27 \pm 2.94) \times 10^{-11}$ | weakly resolved |
| 793b | 10k | $k$ | 7.14 | $6.77 \pm 0.995$ | [6.44, 8.24] |
| | | $\delta$ | 0.208 | $0.190 \pm 0.0419$ | [0.172, 0.256] |
| | | $\beta_N$ | 0.220 | $0.245 \pm 0.0630$ | [0.162, 0.284] |
| | | $\lambda_V$ | 0.120 | $0.124 \pm 0.0129$ | [0.109, 0.130] |
| | | $\mu_V$ | 0.743 | $0.565 \pm 0.127$ | [0.570, 0.885] |
| | | $n_V$ | 0.783 | $0.793 \pm 0.0437$ | [0.777, 0.796] |
| | | $\zeta$ | $1.05 \times 10^{-10}$ | $(2.37 \pm 6.84) \times 10^{-11}$ | weakly resolved |

*Notes.* All numerical values are rounded to three significant figures; scientific notation is retained where fixed decimal rounding would obscure the parameter scale. Bayesian summaries are the relative-posterior-weighted mean and one weighted standard deviation of the accepted SMC particles used in the Bayesian parameter-pair section below. These posterior spread summaries are not formal confidence intervals and should be interpreted separately from the profile-likelihood 95% confidence intervals shown in brackets. Confidence intervals use the likelihood-ratio threshold $\Delta\ell = 1.92$ . The pressure constant $\zeta$ was retained as a nuisance physical/numerical parameter. It remained weakly resolved across datasets and is not interpreted biologically. The resolution labels are local and conditional on the accepted calibration basins; correlated parameter combinations may be constrained even when individual parameters are not independently resolved. For 793b 10k, the reportable 70% necrotic-radius basin is shown rather than the lower-WRMSE but clone-collapsed 80% basin. For 793b 5k, the best reportable/profile-supported value is shown, using the accepted profile extensions where these improved the final likelihood support.

## S8 Model Derivation

This supplementary section contains the full derivation of the phase-field framework summarised in the main-text model implementation section. The purpose of retaining the full derivation here is to preserve the thermodynamic and continuum-mechanics detail supporting the implemented model, while keeping the main text focused on the equations and interpretations most relevant to the confirmation argument.

### S8.1 Model Derivation Details

#### Setup

We start by considering a system of *N* cell populations: *φ*_*i*_ ∈ { *φ*_1_, … , *φ*_*N*_} , as well as two “macro” systems *T* and *H*, corresponding to the total cell population and the host, such that:

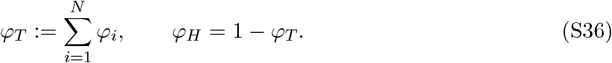

#### General balance law

We assume that these cell populations follow the generalised inhomogeneous continuity equation:

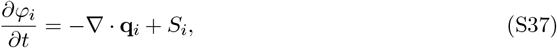

where **q**_*i*_ is the mass flux, and *S*_*i*_ is the inhomogeneous mass source term. The mass flux can be written as **q**_*i*_ = **v**_*i*_*φ*_*i*_, where **v**_*i*_ is the velocity field of cell species *i*.

We write the total flux as **q**_*i*_ = *φ*_*i*_**u**_*s*_ − **J**_*i*_, where **u**_*s*_ is the bulk (mass-averaged) velocity, which we denote the *solid velocity*, and **J**_*i*_ is the diffusive mass flux. The minus sign fixes the convention used in the implemented equations:

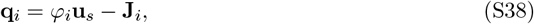

leaving the equations of motion for the cell species fields:

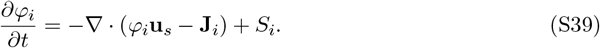

We will now derive the mass flux and solid velocity through thermodynamic principles.

#### S8.1.1 System Free Energy

##### Starting point

We introduce a free energy for each cell population *E*_*i*_, starting from the Helmholtz free energy:

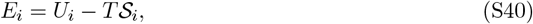

where *T* is the temperature, *U*_*i*_ is the internal energy, and *S*_*i*_ is the entropy of component *i*. We assume an isothermal system.

We assume that *U*_*i*_ arises from the adhesive forces between populations *i* and *j*. The total interaction energy for population *i* is written as:

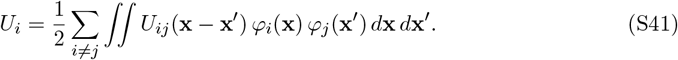

This interaction potential describes the interaction energy as a function of the distance between populations *i* and *j*.

##### Localisation Assumption

We assume that the interaction between cells only occurs when cells are close, and thus that *U*_*ij*_ is localised around **x** and symmetric, *U*_*ij*_ = *U*_*ji*_. Starting from (S41), we localise the system by assuming that *U*_*ij*_ is consistent around any **x**:

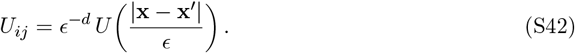

We next make a coordinate change by setting 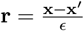, which implies *d***r** = *d***x**^*′*^*/ϵ*^*d*^. Substituting into *U*_*i*_ gives:

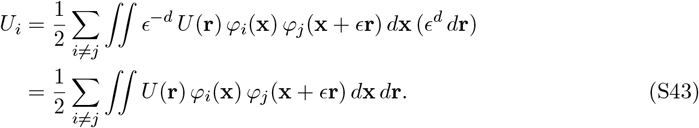

##### Local expansion

Because this describes a local interaction, we can expand *φ*_*j*_ in *ϵ* to remove the dependence on **r**. Using the directional derivative form of a Taylor series:

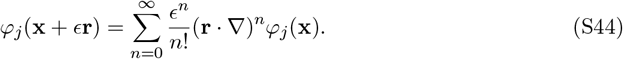

Since 0 *< ϵ* ≪ 1, we retain terms up to *O*(*ϵ*^3^):

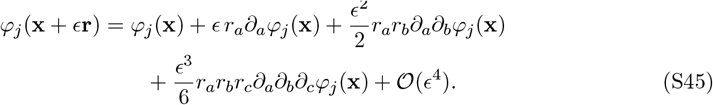

Substituting back into the interaction energy and separating the integral yields:

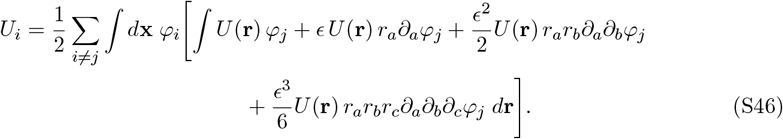

Focusing on the inner integral with respect to **r** gives:

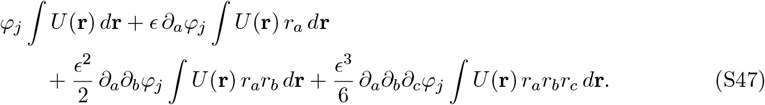

##### Moment reduction under isotropy

We now assume that *U* is isotropic: *U* (**r**) = *U* (|**r**|). As *U* is isotropic (and hence even), the integrals with odd powers of *ϵ* have odd integrands and therefore vanish:

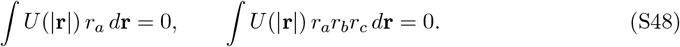

This reduces our expression to the zeroth and second order moments of the energy:

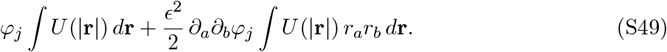

We define the zeroth moment by

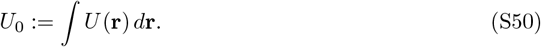

To simplify the second moment, we use a rotation argument and define

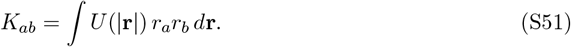

Because *U* is both isotropic and symmetric, it is invariant under rotations of the coordinate system: *U* (|**r**|) = *U* (*R*|**r**|) for *R* ∈ SO(*d*). Applying the change of variables **y** = *R***r**, so that **r** = *R*^*T*^ **y** and *d***r** = *d***y**:

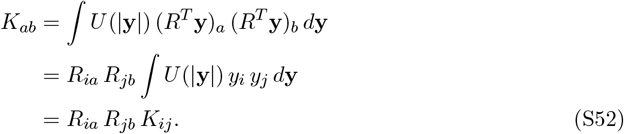

In matrix notation this gives *K* = *R*^*T*^ *KR*, or equivalently *RK* = *KR*. Since *K* commutes with every rotation *R*, the only symmetric matrices with this property are scalar multiples of the identity:

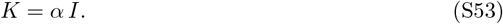

Computing the trace of *K*:

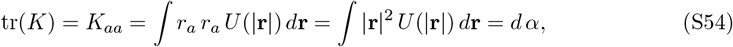

which gives:

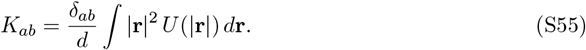

We then define the second moment by

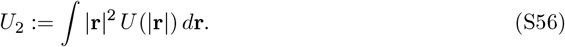

Substituting the zeroth and second order moments back into the interaction energy gives:

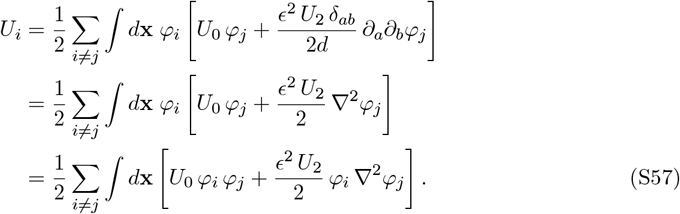

We simplify the Laplacian term through integration by parts:

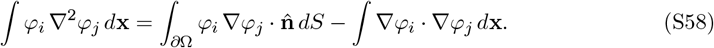

Specifying no-flux boundary conditions for the cell fields 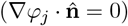, the boundary term vanishes, leaving:

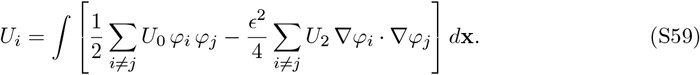

##### Macro-scale reduction

We now split our indices into two macro sets: tumour *T* and host *H*,

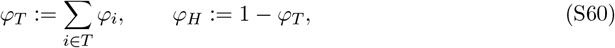

and assume that the cell fields do not interact with the host: 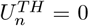. We write individual populations in terms of the total population as *φ*_*i*_ = *w*_*i*_ *φ*_*T*_ , which implies 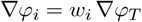.

The remaining terms can now be grouped separately. For the non-gradient contribution:

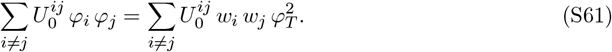

We define

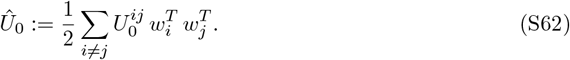

For the gradient contribution:

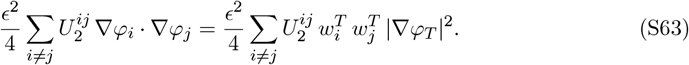

We then define the effective “cell mobility” as

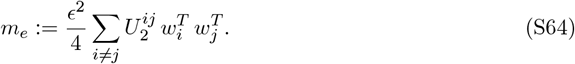

Combining all elements, the interaction energy takes the form:

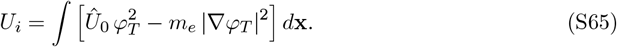

##### Entropy of Mixing

We introduce the entropy of mixing as:

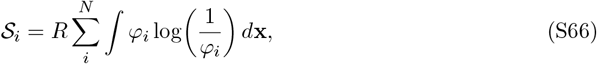

giving us the total free energy:

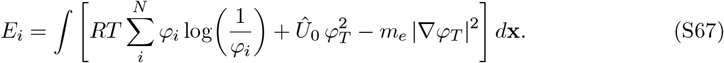

##### Separating bulk and interfacial contributions

Starting from (S67), equivalently written as:

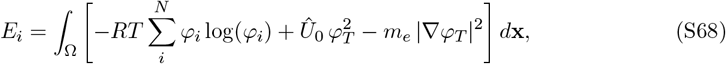

we isolate the first two terms for the Landau expansion, deferring the gradient term:

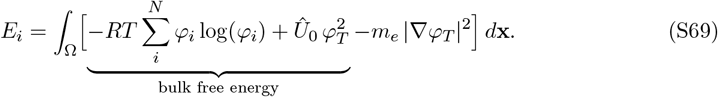

##### Rewriting the entropy term

Recalling the macro definitions *φ*_*T*_ := ∑_*i*_ *φ*_*i*_ and *φ*_*H*_ := 1 − *φ*_*T*_ , we split the sum:

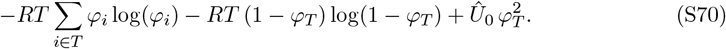

Using the relation *w*_*i*_ = *φ*_*i*_*/φ*_*T*_ , we obtain:

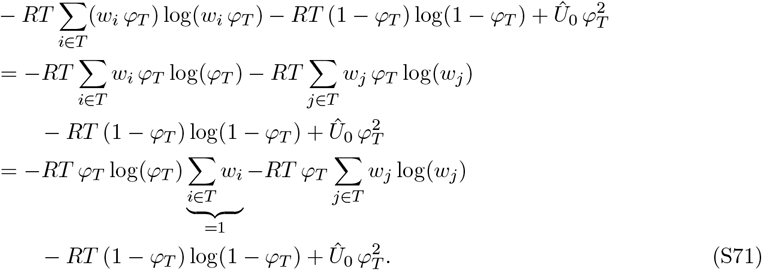

If we assume *w*_1_ = *w*_2_ = = *w*_*N*_ = 1*/N* , then ∑_*j*_∈_*T*_ *w*_*j*_ log(*w*_*j*_) = − log(*N*), so the bulk term becomes

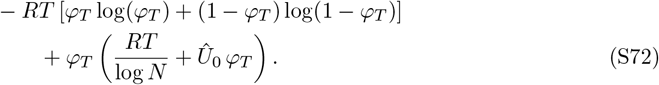

##### Expansion about the symmetric state

We now perform a Taylor expansion of the logarithmic terms around *φ*_*T*_ = 1*/*2. Setting *ψ* = *φ*_*T*_ − 1*/*2:

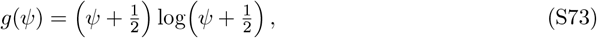

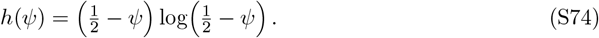

Expanding each around *ψ* = 0:

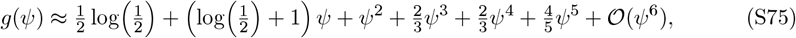

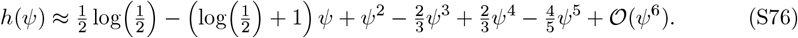

Since *g* + *h* is even, all odd-power terms cancel:

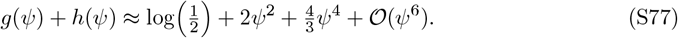

Substituting back and returning to the original coordinate *φ*_*T*_ gives:

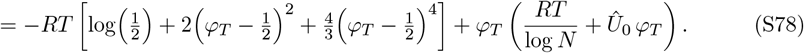

Expanding and collecting like terms yields:

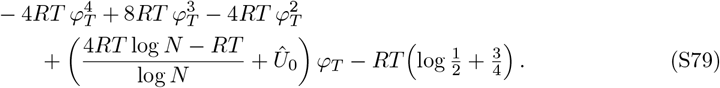

##### Recovering the double-well potential

We now choose

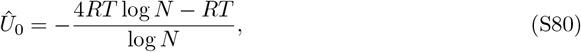

and define

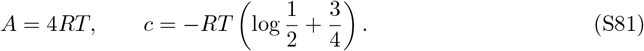

This simplifies the bulk free energy to

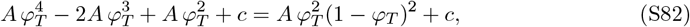

which is the canonical quartic double-well potential, up to a constant *c* that is independent of *φ*_*i*_ and the spatial coordinates (and thus will vanish upon variation).

Performing the Landau expansion, the free energy becomes:

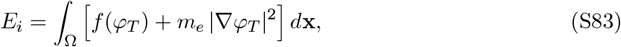

where 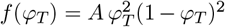, which is of Cahn–Hilliard form. Taking the variational derivative yields the chemical potential *ν*_*T*_ , i.e. the adhesion energy gradient along which cells move:

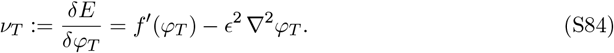

#### S8.1.2 Solid Velocity and Mechanical Closure

We now derive the form of the solid velocity **u**_*s*_. Starting from the free energy functional:

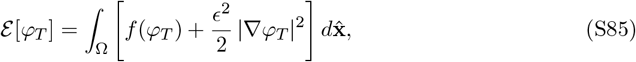

we differentiate with respect to time:

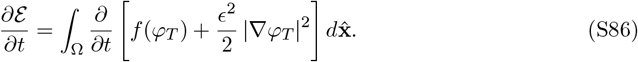

We separate the two terms under the integral. For the first:

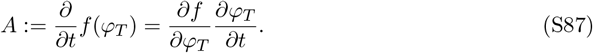

For the second:

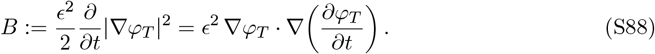

Using the identity ∇ · (*a* ∇*b*) = ∇*a* · ∇*b* + *a* ∇^2^*b* with *a* = *∂*_*t*_*φ*_*T*_ and *b* = *φ*_*T*_ :

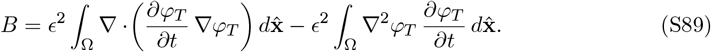

Assuming no-flux boundary conditions 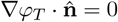, we apply the divergence theorem:

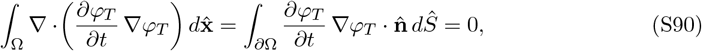

so that *B* reduces to:

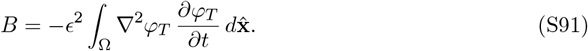

Reassembling and factoring out *∂*_*t*_*φ*_*T*_ :

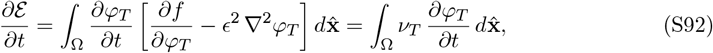

where we have identified the chemical potential 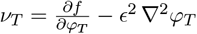.

Substituting the equations of motion (S39):

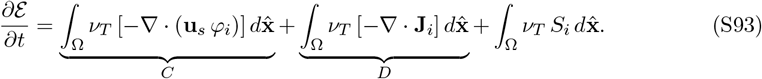

##### Analysis of C

Expanding the divergence:

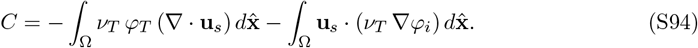

Applying the incompressibility constraint ∇ · **u**_*s*_ = *S*_*T*_ :

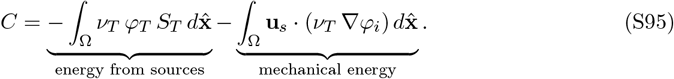

The mechanical closure used in the implemented model is Darcy-like. We therefore write:

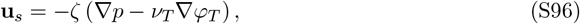

where *p* is the effective mechanical pressure and *ζ* is the pressure-relaxation constant. Taking the divergence and imposing ∇ · **u**_*s*_ = *S*_*T*_ gives:

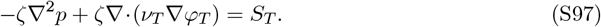

Rearranging, we obtain a Poisson equation for the mechanical pressure *p*:

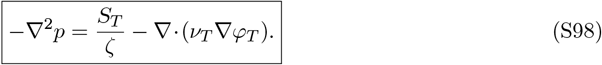

When this equation is satisfied, the volume-balance constraint holds, and the solid velocity is:

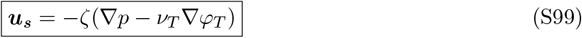

##### Analysis of D

Integrating by parts:

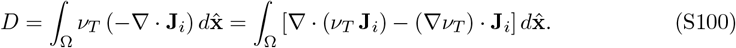

Using the divergence theorem and boundary conditions, the first term vanishes:

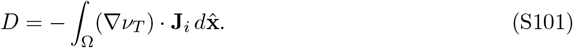

Since this term contains no source, we again invoke the dissipation argument: (∇*ν*_*T*_) ·**J**_*i*_ ≥ 0, so that **J**_*i*_ and ∇*ν*_*T*_ are aligned. Assuming the system maximises energy dissipation, we obtain the constitutive relation for the mass flux:

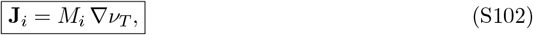

with *M*_*i*_ = *m*_*i*_*φ*_*i*_ for the population-specific mobility used in the main text.

#### S8.1.3 Nutrient Field

We denote the nutrient concentration field by *n*(**x**, *t*). At the continuum scale, we assume that nutrient transport is governed by diffusion together with local cellular consumption, giving the general reaction-diffusion balance

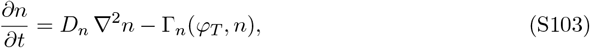

where *D*_*n*_ is the nutrient diffusivity and Γ_*n*_ is the net nutrient uptake term.

The key modelling assumption is that nutrient diffusion occurs on a much faster timescale than tumour growth. In particular, nutrient redistribution takes place on the order of minutes, whereas spheroid growth and compositional changes occur over hours to days. This separation of timescales implies that, on the timescale over which *φ*_*T*_ evolves, the nutrient field rapidly relaxes to equilibrium.

Under this quasi-steady assumption, the transient term is negligible relative to diffusion and consumption, so that

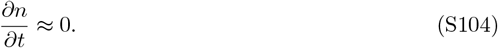

The nutrient balance therefore reduces to the steady-state reaction-diffusion equation

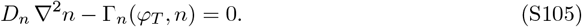

This provides the form used throughout the model: at each tumour configuration, the nutrient field is treated as being in instantaneous equilibrium with the current cell distribution, rather than evolving dynamically on the same timescale as the tumour phase fields.

For the viable–necrotic model used here, the uptake term is Γ_*n*_ = *u*_*V*_ *φ*_*V*_ *n* with necrotic uptake omitted. The corresponding quasi-steady equation is therefore ∇ · (*D*_eff_ ∇*n*) − *u*_*V*_ *φ*_*V*_ *n* = 0, which gives the exponential penetration-depth scale used in the nondimensionalisation section above.

#### S8.1.4 Source Terms

All biochemical processes, including cell proliferation, death, and phenotypic conversion, are encoded in the source terms *S*_*i*_. At this stage of the project we do not consider broad phenotypic conversion pathways such as carcinogenesis in the viable–necrotic model, other than the transfer of viable cells into the necrotic compartment through death. However, we do provide a general framework for integrating such pathways in Section S1.1. We decompose each *S*_*i*_ into three conceptually distinct components: incoming mass *I*_*i*_, outgoing mass *O*_*i*_, and mass-transfer terms *T*_*i*_:

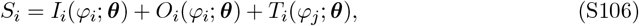

where each function depends on the parameter set ***θ***.

At the general level, this decomposition separates net production, net loss, and inter-population exchange without yet committing to a particular biological mechanism. The specific functional choices used in the viable–necrotic model are introduced only after the general transfer and vector formulations have been established.

#### S8.1.5 Phenotypic Conversion

The previous expressions define the local gain and loss processes for a single population. To extend the framework to multiple interacting phenotypes, we now introduce two distinct forms of mass transfer between populations. The first redistributes newly generated mass. Let *I*_*i*_ denote the total incoming mass generated by population *i*, and let *p*_*i*_ denote the fraction of that incoming mass that remains in population *i*. The remaining fraction (1 − *p*_*i*_) is transferred to another population *j*. This yields

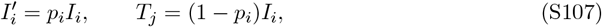

so that 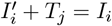 and mass is conserved.

The second form of transfer acts directly on an existing population, independently of whether new mass is being created. If a fraction or rate of population *i* converts into population *j*, we may write this as

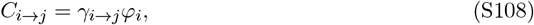

where *γ*_*i*→*j*_ denotes the conversion rate. This contributes a loss term −*C*_*i*→*j*_ to population *i* and an equal gain term +*C*_*i*→*j*_ to population *j*, so that the transfer remains mass conservative.

##### Generalised Transfer

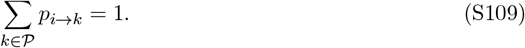

Incoming mass for population *i* then becomes 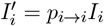 and each recipient population *k* ≠ *I* obtains a transfer term *T*_*k*_ = *p*_*i*→*k*_*I*_*i*_.

Likewise, for direct population-to-population conversion we introduce rates *γ*_*i*→*k*_ for *i* ≠ *k*, and collect the associated net conversion terms into a conversion operator acting on the state vector. This provides a general bookkeeping structure for both growth-mediated redistribution and direct inter-population exchange before introducing the full vector form used in implementation.

#### S8.1.6 Vector Formulation

To streamline implementation and generalisation, while also unlocking hardware-accelerated linear-algebra kernels, we collect all *M* cell-phase fields into the state vector

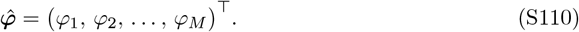

We fix the necrotic population to index *M* , so that *φ*_*M*_ := *φ*_*N*_. We then define likewise-stacked mass fluxes

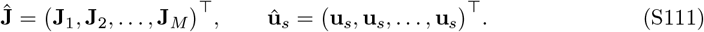

Equation (S39) may then be written compactly as

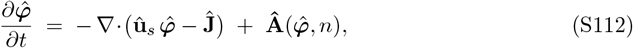

where **Â** is the generalised source vector whose entries encode phenotype-specific proliferation, death, and conversion rates.

To construct **Â**, we first define the transfer matrix **P**, whose entries specify how newly generated mass from source population *i* is distributed across recipient populations *k*:

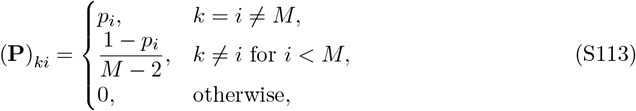

so that each column sums to one, 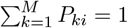.

Next, we define a direct-conversion matrix **C**, whose off-diagonal entries encode conversion of existing mass from population *i* into population *k*, while the diagonal terms collect the net loss from each source population:

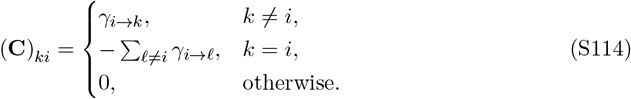

By construction, each column of **C** sums to zero, so direct conversion redistributes mass between populations without net creation or loss. We then define the production vector 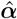, whose entries are the net production rates *α*_*i*_ generated by each population. The corresponding loss and transfer-to-necrosis effects are collected in the death matrix **D**:

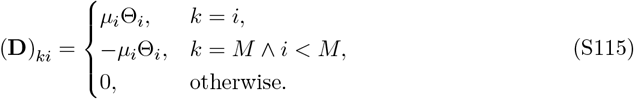

The general source vector is therefore

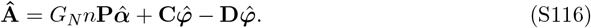

#### S8.1.7 Specialisation to the V–N Model

We now recover the viable–necrotic model used in this project as the simplest specialisation of the general framework above. We restrict cell phenotypes to a duet of viable cells, which may include tumour cells, stem-like cells, and other proliferative states, and necrotic cells. In this setting we do not include broader inter-phenotype transfer pathways, and only resolve nutrient-dependent growth, viable-cell death into the necrotic compartment, and necrotic clearance.

##### Nutrient-Dependent Proliferation

Growth of the viable population is taken to be proportional to both nutrient availability and viable mass, so that the incoming term is

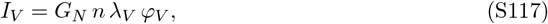

where *λ*_*V*_ is the effective viable-cell proliferation rate, *n* is the local nutrient concentration, and *G*_*N*_ is a global necrosis-mediated inhibition factor defined below.

##### Nutrient-Dependent Death

Viable-cell death is encoded through the outgoing term

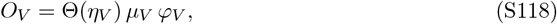

where *µ*_*V*_ is the viable-cell death rate. The switch Θ is taken as the smooth nutrient-response function

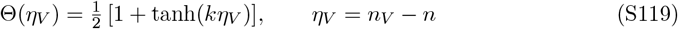

which interpolates between survival when nutrient levels exceed the threshold *n*_*V*_ and death under nutrient depletion. The parameter *k* controls the steepness of the transition, and the smooth hyperbolic tangent is used in place of a discontinuous Heaviside switch.

##### Global Necrotic Feedback

To reproduce the late-stage, approximately logistic growth behaviour observed in spheroid systems, we introduce the global necrotic feedback factor

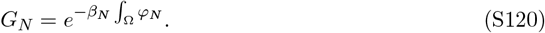

Here *β*_*N*_ controls the strength of the feedback. Rather than representing a single biological mechanism, *G*_*N*_ is interpreted as an effective negative feedback that captures the aggregate influence of necrosis-associated effects that suppress further expansion. These may include increased mechanical stress, reduced tissue mobility arising from necrotic regions, with necrotic cells assumed immotile, as well as the release of necrosis-associated factors that locally inhibit proliferation.

The source terms therefore reduce to

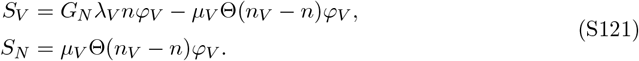

Here necrotic material accumulates from the death of the viable population; clearance is omitted by setting *µ*_*N*_ = 0. In vector form this corresponds to

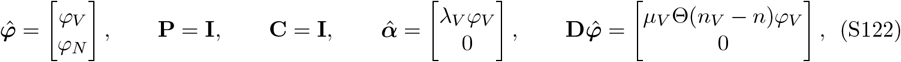

with the source vector

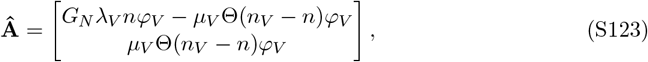

and hence

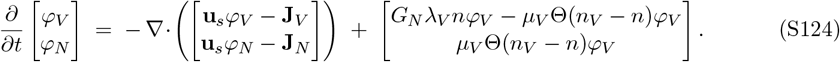

The source terms in the viable–necrotic model therefore depend on the parameter set

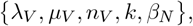

along with the evolving nutrient field *n* and phase fields *φ*_*V*_ , *φ*_*N*_. In summary, the thermodynamic derivation defines the transport structure, the general source-term framework defines how production, loss, and transfer are organised, and the viable–necrotic model specifies the particular biological mechanisms currently resolved by the project.

### S8.2 Drug-Response Source-Term Perturbations

Drugged simulations were implemented by introducing a relative drug-concentration field *d*(**x**, *t*), with *d* = 0 corresponding to no local drug exposure and *d* = 1 corresponding to the imposed boundary exposure. The effect of the drug on each viable population *i* is mediated by a bounded activity function

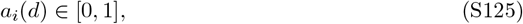

which maps local drug concentration to local pharmacological activity. In the simulations shown here, we used the Hill-type activity function

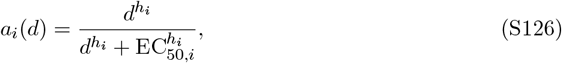

unless otherwise stated. Here *h*_*i*_ is the Hill coefficient and EC_50,*i*_ is the relative drug concentration producing half-maximal activity.

For a viable population *i*, the untreated source terms depend on the proliferation rate *λ*_*i*_, death rate *µ*_*i*_, and nutrient survival threshold *n*_*i*_. Drug action was introduced by locally perturbing these quantities through three independently enabled mechanisms.

#### Growth Suppression

Growth suppression reduces the effective proliferation rate of the viable population:

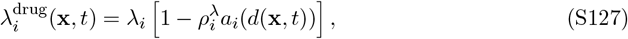

where 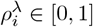 sets the maximum fractional reduction in proliferation. Thus, when *a*_*i*_(*d*) = 0, the untreated proliferation rate is recovered, while when *a*_*i*_(*d*) = 1, proliferation is reduced by the fraction 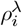.

#### Direct Drug-Induced Death

Direct cytotoxicity is represented as an additional viable-to-necrotic transfer,

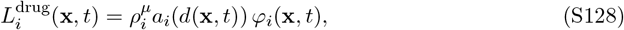

where 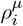 is the maximum drug-induced death rate and *φ*_*i*_ is the local volume fraction of viable population *i*. This modifies the viable and necrotic source terms according to

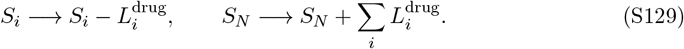

Direct drug-induced death therefore conserves local cellular material by transferring viable density into the necrotic compartment.

#### Nutrient Sensitisation

Nutrient sensitisation increases the nutrient concentration required for survival. Consistent with the death-activation convention in Equation (S119), the drug-shifted nutrient threshold is

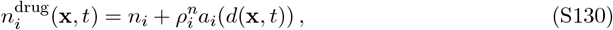

where 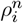 is the maximum drug-induced threshold shift. The corresponding drug-modified nutrient-death switch is

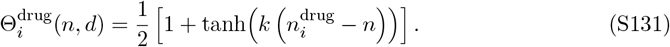

Increasing 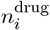 therefore expands the region in which cells experience nutrient-mediated death. Combining the three mechanisms, the drug-modified source terms for viable populations and the necrotic compartment may be written as

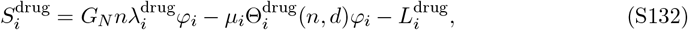

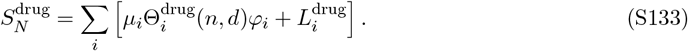

Any subset of growth suppression, direct drug-induced death, and nutrient sensitisation can be enabled independently. Setting 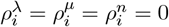 recovers the untreated source terms.

## S9 Bayesian Parameter-Pair Distributions

The following plots show the pairwise parameter distributions for the accepted Bayesian calibration samples from each Browning dataset and initial seeding density.

**Figure S1:**
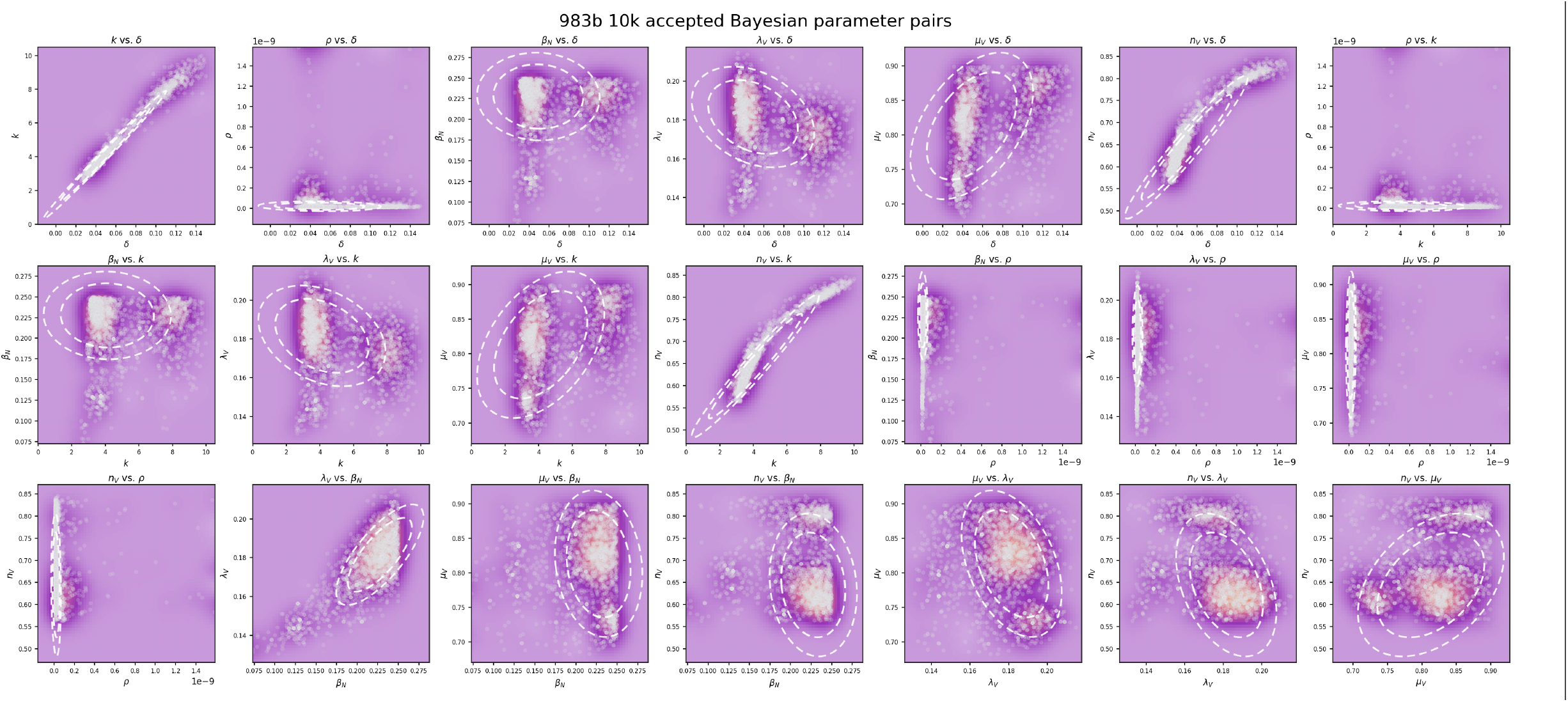
Bayesian parameter-pair distributions for accepted calibration samples from the Browning WM983b 10k dataset.

**Figure S2:**
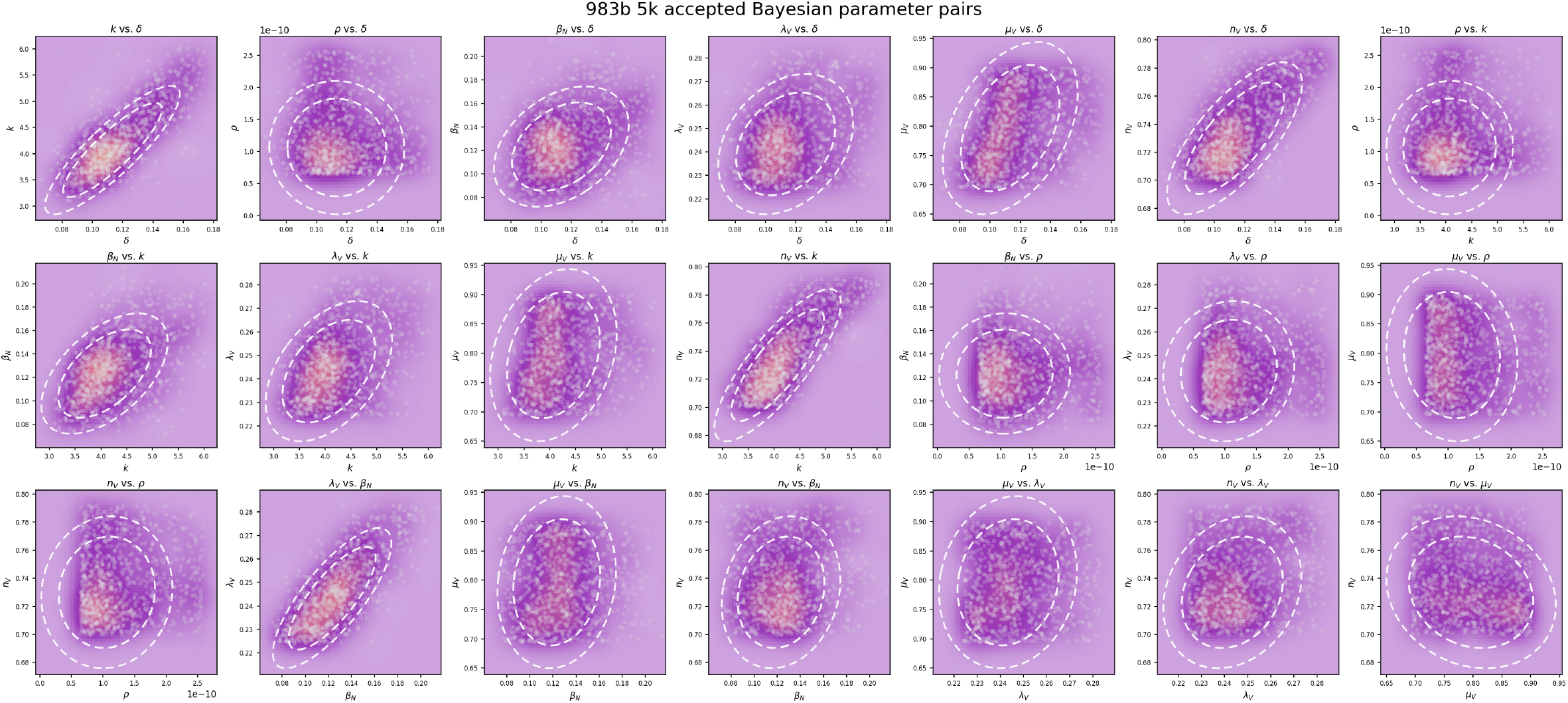
Bayesian parameter-pair distributions for accepted calibration samples from the Browning WM983b 5k dataset.

**Figure S3:**
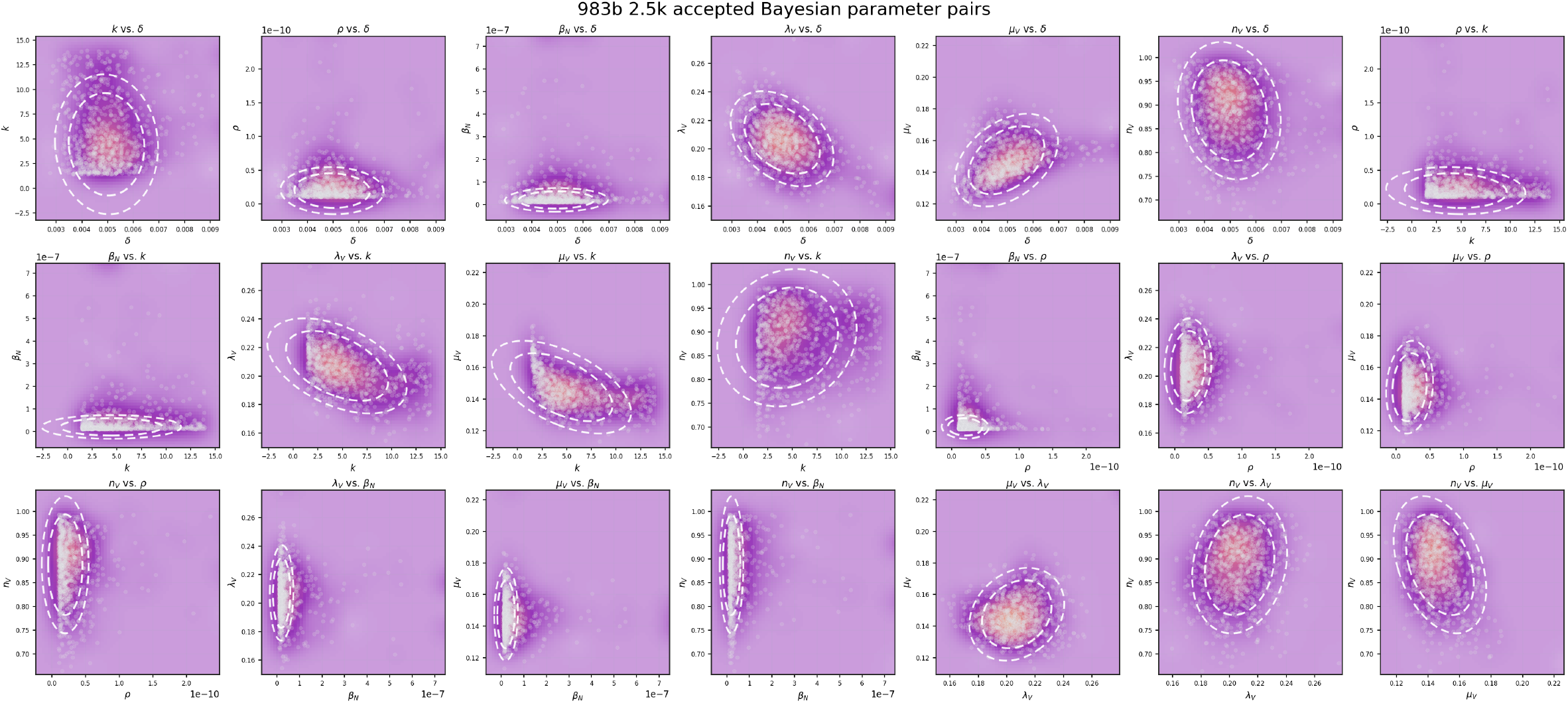
Bayesian parameter-pair distributions for accepted calibration samples from the Browning WM983b 2.5k dataset.

**Figure S4:**
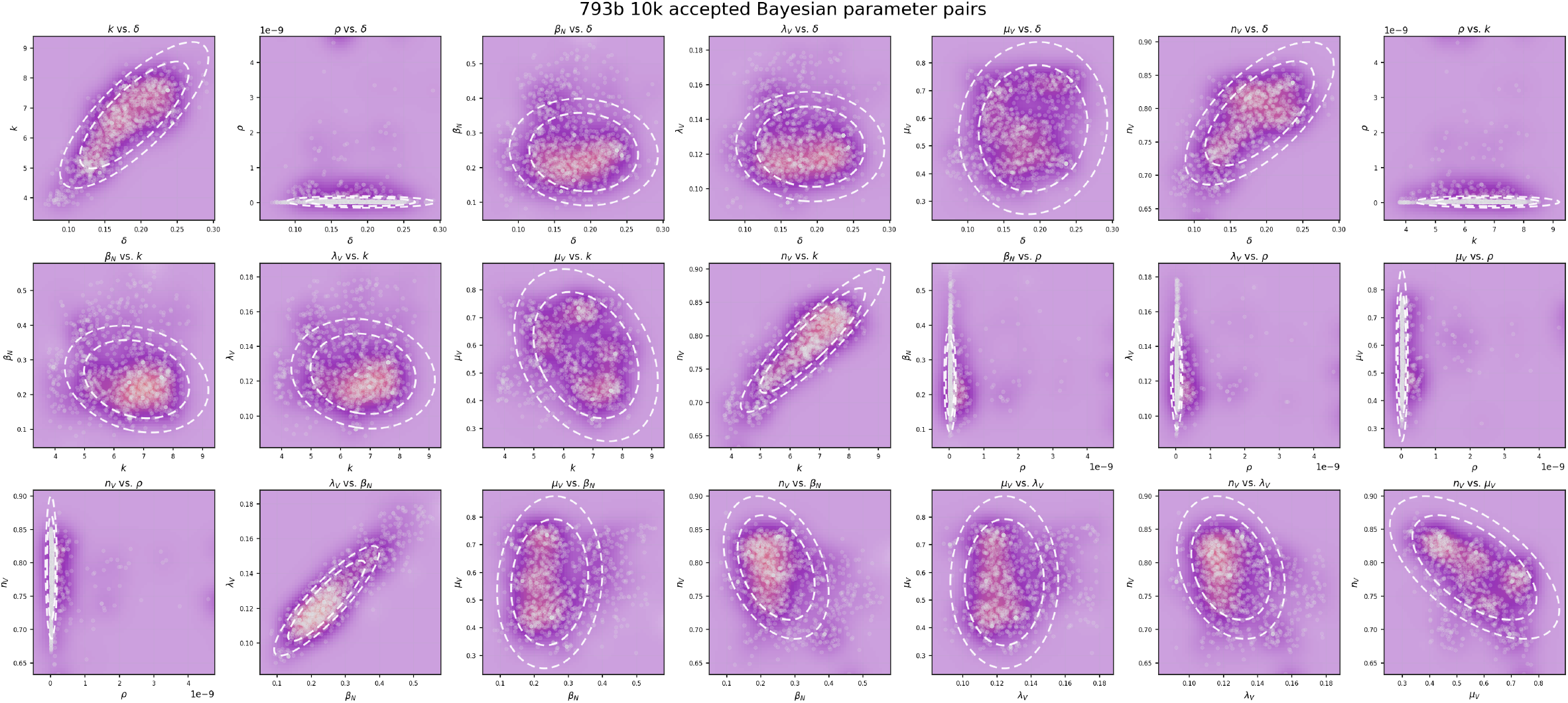
Bayesian parameter-pair distributions for accepted calibration samples from the Browning WM793b 10k dataset.

**Figure S5:**
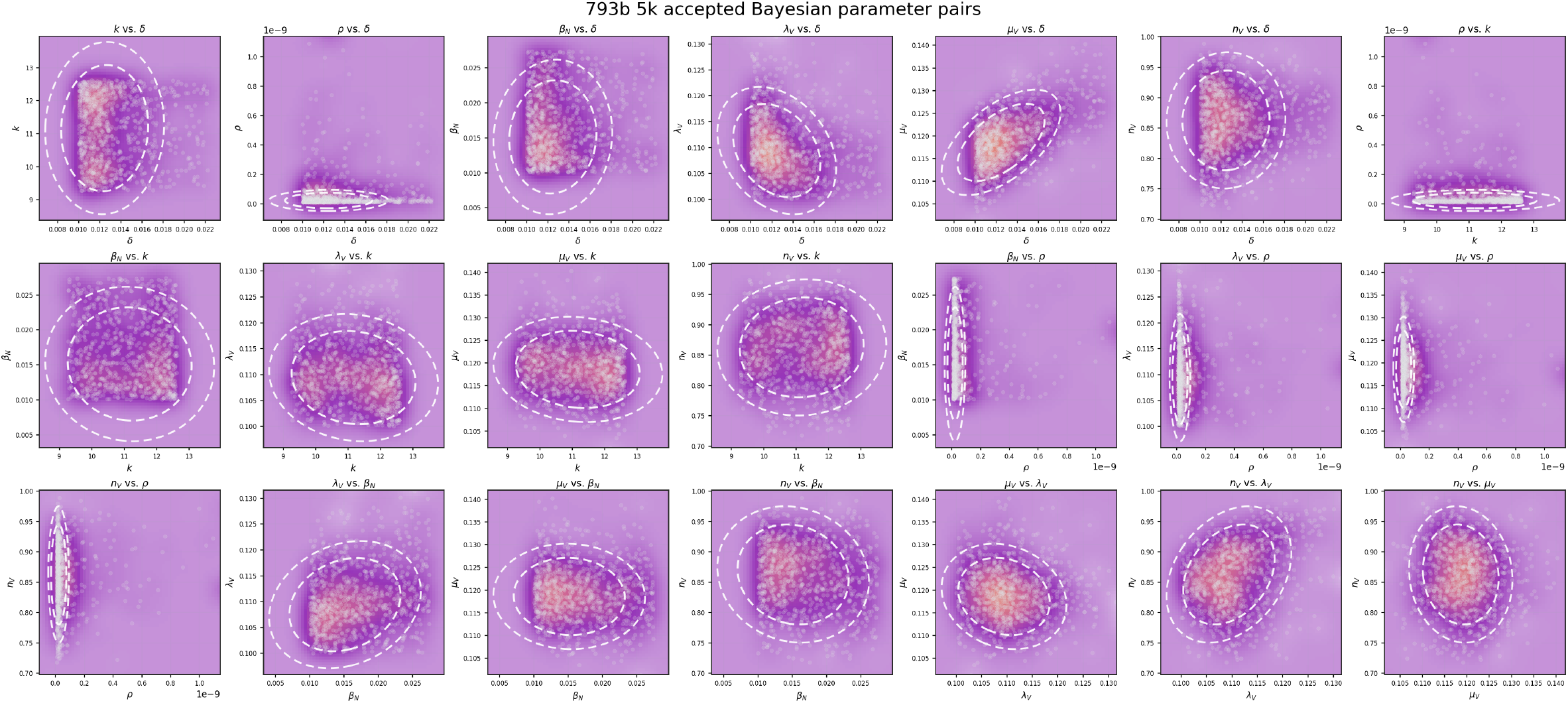
Bayesian parameter-pair distributions for accepted calibration samples from the Browning WM793b 5k dataset.

**Figure S6:**
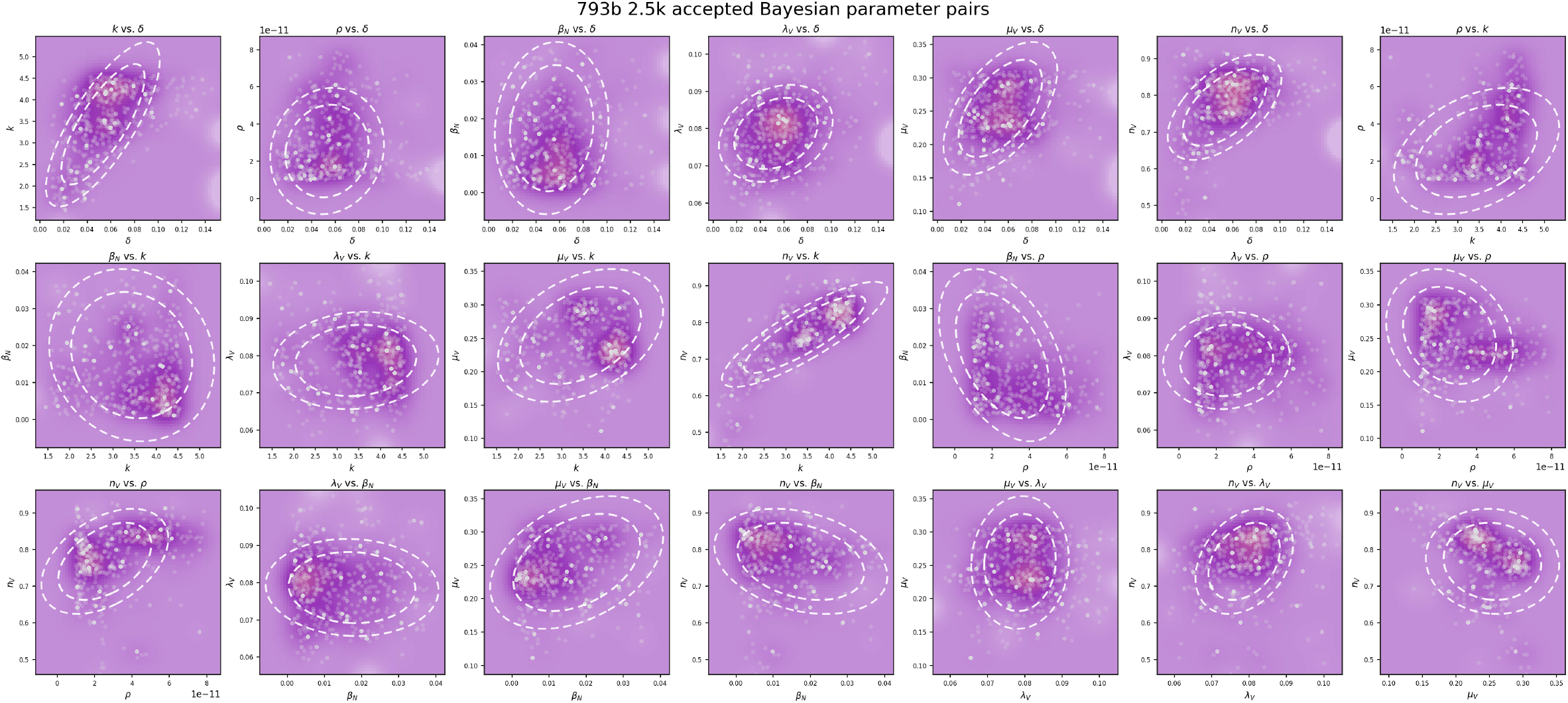
Bayesian parameter-pair distributions for accepted calibration samples from the Browning WM793b 2.5k dataset.

